# Pseudorabies virus hijacks DDX3X, initiating an addictive “mad itch” and immune suppression, to facilitate viral spread

**DOI:** 10.1101/2023.05.09.539956

**Authors:** Shane J. F. Cronin, Miguel A. Tejada, Ren Song, Kathlyn Laval, Domagoj Cikes, Ming Ji, Annalaura Brai, Johannes Stadlmann, Maria Novatchikova, Thomas Perlot, Omar Hasan Ali, Lorenzo Botta, Thomas Decker, Jelena Lazovic, Astrid Hagelkruys, Lynn Enquist, Shuan Rao, Orkide O. Koyuncu, Josef M. Penninger

## Abstract

Infections with defined Herpesviruses, such as Pseudorabies virus (PRV) and Varicella zoster virus (VZV) can cause neuropathic itch, referred to as “mad itch” in multiple species. The underlying mechanisms involved in neuropathic “mad itch” are poorly understood. Here, we show that PRV infections hijack the RNA helicase DDX3X in sensory neurons to facilitate anterograde transport of the virus along axons. PRV induces re-localization of DDX3X from the cell body to the axons which ultimately leads to death of the infected sensory neurons. Inducible genetic ablation of *Ddx3x* in sensory neurons results in neuronal death and “mad itch” in mice. This neuropathic “mad itch” is propagated through activation of the opioid system making the animals “addicted to itch”. Moreover, we show that PRV co-opts and diverts T cell development in the thymus via a sensory neuron-IL-6-hypothalamus-corticosterone stress pathway. Our data reveal how PRV, through regulation of DDX3X in sensory neurons, travels along axons and triggers neuropathic itch and immune deviations to initiate pathophysiological programs which facilitate its spread to enhance infectivity.

## Introduction

Pruritus (or itch) is a normal protective physiological response caused by the stimulation of itch-sensing C-fiber nerve endings which then elicit the behavioral response of scratching or rubbing to remove the irritant. Under certain pathophysiological conditions without apparent pruritogenic stimuli, itch, like pain, can become chronic, a condition called neuropathic itch (Oaklander, 2011). A central feature of neuropathic itch is uncontrollable scratching into deep tissues which is almost always accompanied by loss of pain sensation that can result in severe self-injury (Oaklander, 2011). Causes of neuropathic itch generally involve damage to sensory peripheral nerves which innervate the face such as in multiple sclerosis, an autoimmune disease involving the myelin sheathes of nerves, or in surgical nerve destruction to treat trigeminal neuralgia (Fitzek et al., 2006; McVeigh et al., 2018; Otters et al., 2014). Very little is known about the cellular and molecular mechanisms underpinning neuropathic itch and various medications such as anti-histamines, corticosteroids and pain medications are largely ineffective. Unlike other itch-related pathologies which arise from immune dysregulation such as psoriasis or atopic dermatitis, and for which numerous model systems exist, there is a scarcity of animal models that reflect the clinical manifestations of neuropathic itch. Hence, most of what is known about neuropathic itch is based on sporadic human case reports.

Pseudorabies virus (PRV, or Suid Herpesvirus 1 (SuHV-1)) is an alpha herpesvirus (α-HV, genus Varicellovirus) swine pathogen and is closely related to the human α-HVs herpes simplex virus type 1 and 2 (HSV1 and HSV2) and varicella-zoster virus (VZV), the chickenpox-causing virus (Mettenleiter, 2000). PRV infections start in the nasal epithelium from where progeny viral particles enter nerve endings that innervate the nasal mucosa and spread retrogradely to infect peripheral nerve system (PNS) neurons where the virus can remain latent. Upon reactivation in sensory neurons, PRV spreads along axons anterogradely from neuron to neuron and eventually to epithelial cells to allow spread to another host (Babic et al., 1994; Damann et al., 2006; Field and Hill, 1974; Koyuncu et al., 2020). In non-natural (non-swine) hosts, PRV infection is known to cause severe acute neuropathy. The main characteristic symptoms observed in all non-natural PRV-infected animals such as domestic cats, dogs, cattle, sheep, or goats and wild animals (e.g. rodents, raccoons, rabbits) is the prevalence of a “mad itch” - severe pruritus in the head and neck areas accompanied by self-mutilation (Laval and Enquist, 2020). The underlying mechanisms of PRV-associated “mad itch” remain poorly understood.

Here, we uncover novel pathophysiological mechanisms involved in “mad itch” providing insights into critical processes that govern viral-neuronal interactions and neuropathic itch.

## Results

### PRV infection of neurons reduces DDX3X levels

During PRV infection of both natural and non-natural hosts, PRV preferentially infects neurons in dorsal root ganglia (DRG), trigeminal ganglia (TG), and superior cervical ganglia (SCG). After replication in neuronal cell bodies, the progeny virus particles can spread retrogradely or anterogradely along axons causing neuronal death and, in non-natural hosts, leading to “mad itch”. Previously, we uncovered host proteins which associate with PRV particles and identified DDX3X in PRV virus particles (Kramer et al., 2011). DDX3X belongs to the DEAD-box RNA helicase superfamily 2 that has widespread functions in RNA metabolism, including transcription, RNA processing, splicing, decay and translation (Fairman-Williams et al., 2010; Fuller-Pace, 2006; Linder and Jankowsky, 2011). DDX3X has also been implicated in regulating infections from a host of viruses including SARS-COV-2 (Ciccosanti et al., 2021), Zika (Nelson et al., 2021), influenza (Kesavardhana et al., 2021), West Nile (Brai et al., 2019), HSV1 (Khadivjam et al., 2017), and herpesvirus (Kramer et al., 2011; Loret et al., 2008; Varnum et al., 2004), without clear mechanistic understandings on its role.

To assess a potential mechanistic role for DDX3X in PRV infections, we first investigated DDX3X protein levels in cultured mouse primary SCG neurons infected with PRV and found that DDX3X protein levels were reduced in a time-dependent manner after infection (**Figure 1a,b**). By 24 hours, DDX3X levels were diminished from baseline by more than 60% compared to mock treatment. These data were validated by immunofluorescence staining in which cell body-localized (predominantly peri-nuclear localized) DDX3X levels were dramatically decreased after PRV infections (**Figure 1c; Supplemental Figure S1a,** hereafter referred to as **Figure S1a**). PRV infections also reduced DDX3X levels in cultured mouse DRG neurons (**Figure S1b-d**). Thus, PRV infections result in downregulation of DDX3X protein levels in mouse sensory neurons.

**Figure 1.**
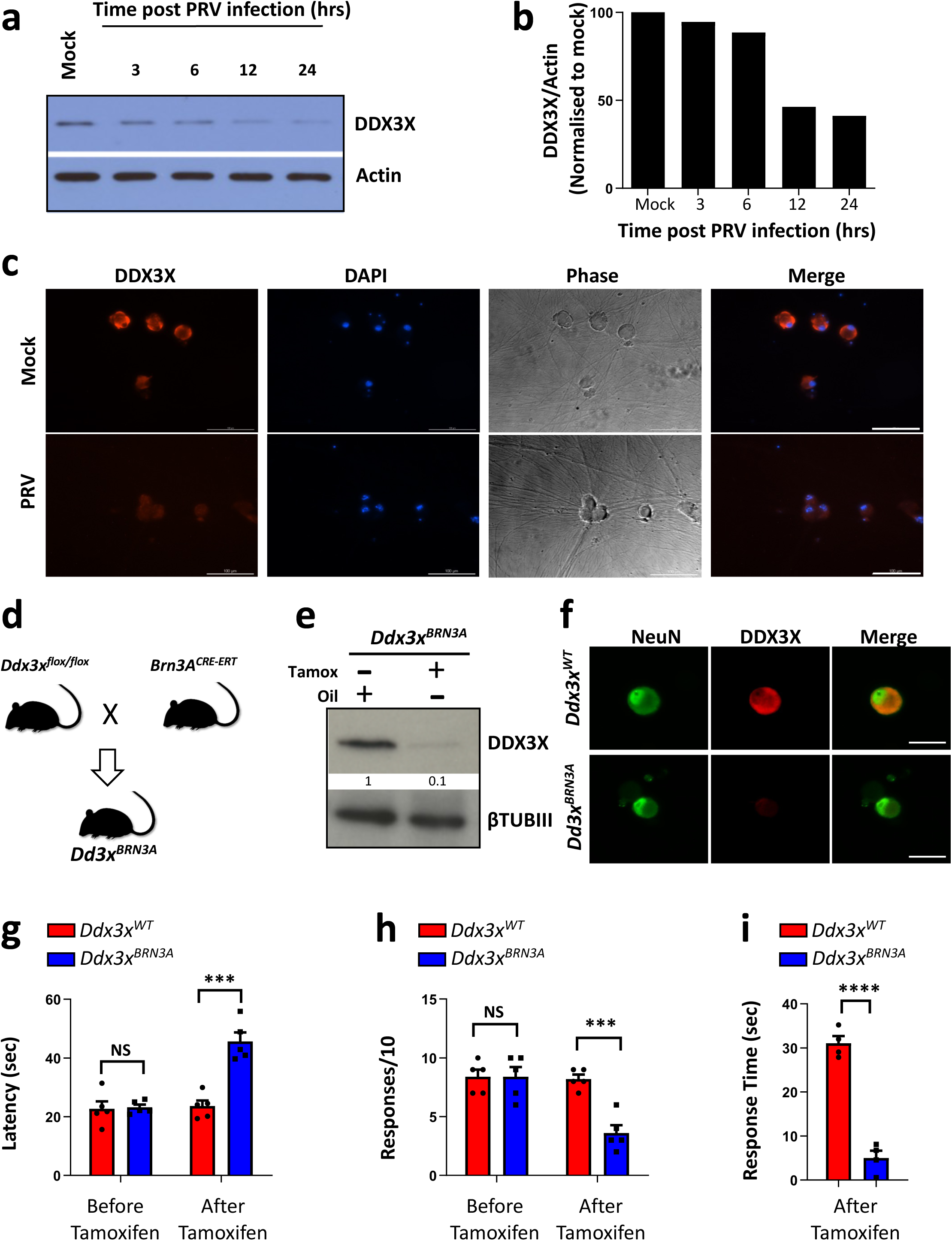
Pseudorabies virus (PRV) infection reduces DDX3X levels in neurons. **a,b,** Western blot time course of superior cervical ganglion (SCG) neurons mock infected and infected with PRV (MOI = 10) and blotted for DDX3X protein expression 3, 6, 12 and 24 hours post infection. Actin is used as a loading control (**a**). Quantification of DDX3X protein levels normalized to mock treatment (no virus infection) (**b**). **c,** Representative immunofluorescence images of DDX3X in cultured SCG neurons mock infected and infected with PRV for 12 hours. DAPI staining and phase contrast images are also shown. Scale bar, 100µm. **d,** Schematic depicting the breeding to temporally ablate *Ddx3x* specifically in small-diameter DRG sensory neurons with tamoxifen treatment (2mg daily for 5 consecutive days) using the *Brn3A-CreERT* driver line. **e,** Western blot of DDX3X from cultured DRG neurons of *Ddx3x^BRN3A^* mice vehicle treated (oil) or 144 hours after initial tamoxifen (tamox) treatment. **f**, Representative immunofluorescence images of cultured DRG sensory neurons from *Ddx3x^BRN3A^*mice vehicle- treated (oil) or 144 hours after initial tamoxifen (tamox) treatment. Beta-tubulin-III (βTUBIII) is used as a loading control. Anti-NeuN is used to mark neuronal nuclei. Scale bar, 50µm. **g,** Contact heat withdrawal latencies using the hotplate assay (52°C) for control *Ddx3x^WT^* (N=5) and littermate *Ddx3x^BRN3A^*(N=5) female mice before and two weeks after tamoxifen treatment. Individual mice for each genotype are shown. **h**, Pin-prick assay on control *Ddx3x^WT^* (N=5) and littermate *Ddx3x^BRN3A^* (N=5) female mice before and two weeks after tamoxifen treatment. Individual mice for each genotype are shown. **i,** Nocifensive responses in 3-minute time frames after intraplantar injection of 2 μg of capsaicin into the left hind paw of control *Ddx3x^WT^* (N=4) and littermate *Ddx3x^BRN3A^*(N=4) female mice two weeks after tamoxifen treatment. The experiment was repeated two additional times with similar results. Individual mice for each genotype are shown (g,h,i). Data are shown as means ± s.e.m. Multiple t-test (g,h); two-tailed unpaired Student’s t test (i). * **P < 0.001; ****P < 0.0001; NS, not significant.

### *Ddx3x* deletion in peripheral sensory neurons induces “mad itch” and self-mutilation

As PRV infection manifests the “mad itch” during infection of adult animals and neuropathic itch in shingles primarily affects older people we hypothesized that ablation of *Ddx3x* in adult mice might cause a neuropathic itch phenotype. To genetically ablate *Ddx3x* in adult sensory neurons, we crossed our *Ddx3x* floxed (*Ddx3x^flox/flox^*) mouse (Szappanos et al., 2018), in which exon 2 of the gene is floxed by loxP recombination sites, to the tamoxifen-inducible, sensory neuronal *Cre*-driver line *Brn3A-Cre-Ert* (O’Donovan et al., 2014) to generate *Ddx3x^BRN3A^* mice (**Figure 1d**). Using a *TdTomato* reporter showed that this *Cre*-expressing line targets primarily the small-diameter CGRP-positive and IB4-binding DRG neurons (**Figure S2**). When crossed to a *YFP^floxSTOPflox^* reporter line, one week after tamoxifen treatment, the small-diameter DRG neurons were YFP-positive (**Figure S3a,b**). When crossed to *Ddx3x^flox/flox^*mice, tamoxifen treatment induced DDX3X ablation (**Figure 1e,f**). Interestingly, 3 weeks after tamoxifen treatment *Ddx3x^BRN3A^*mice displayed significantly reduced pain-like behaviors compared to littermate *Ddx3x^WT^* controls as well as compared to pre-treated *Ddx3x^BRN3A^* mice (**Figure 1g- i**).

Strikingly, deletion of *Ddx3x* in small-diameter DRG neurons of adult mice caused significant head and body lesions in female *Ddx3x^BRN3A^*mice (**Figure 2a**). These lesions were a result of self-mutilation as they also manifested in tamoxifen-treated, singly housed *Ddx3x^BRN3A^* female mice. These mice demonstrated intense behaviors which we collectively herein refer to as “mad itch” including facial and nape scratching and biting of the flank regions, as well as incessant head shakes, and facial and nape paw swipes – with these behaviors culminating in the appearance of self-inflicted severe skin lesions (**Figure 2a,b**). Indeed, the self-mutilation behavior could become so intense as to penetrate deep into the tissue, reminiscent of neuropathic itch seen in human patients (Oaklander, 2011). When observed by independent investigators, these *Ddx3x^BRN3A^* female mice displayed near-constant “mad itch”-like behaviors (**Figure 2c,d; Supplementary Videos 1 and 2**). The skin lesions became apparent in *Ddx3x^BRN3A^*female mice approximately 4 weeks after tamoxifen treatment with a 100% incidence, while vehicle treatment resulted in no such self-inflicted lesions (**Figure 2e**). As expected, inducible *Ddx3x* deletion resulted in loss of the major small-diameter C-fiber neuronal subtypes (CGRP-positive, IB4-binding, and tyrosine hydroxylase-positive (TH+)) while preserving the larger diameter, *Cre*-negative myelinated A-fibers as detected by Parv (marker for proprioceptors) and NF200 (marker for Aβ fibers) (**Figure 2f,g; Figure S3c-f**). Thus, deletion of *Ddx3x* in small-diameter, peripheral sensory neurons resulted in neuronal death and, in all animals with 100% penetrance, in intractable and self-mutilating “mad itch”.

**Figure 2.**
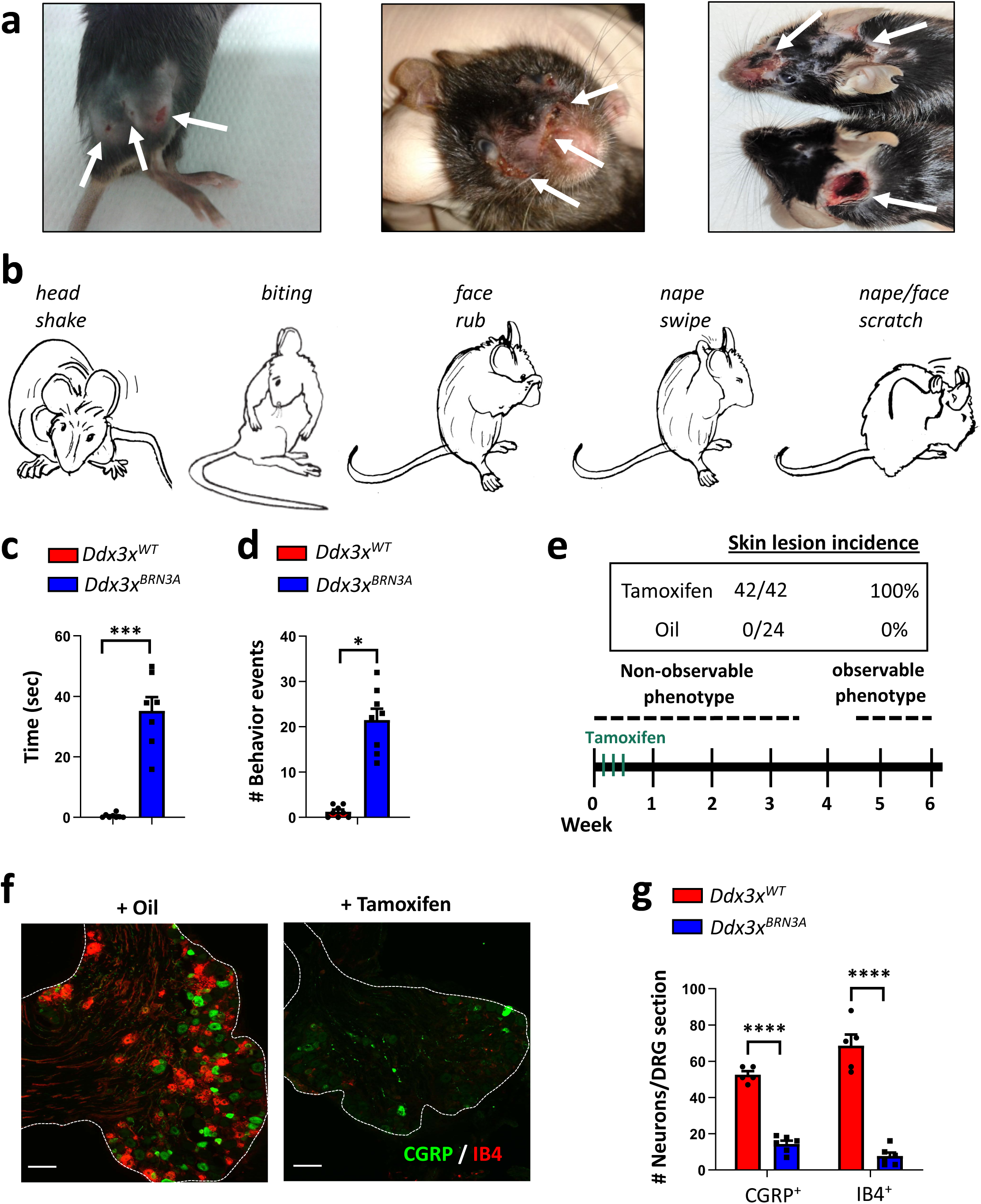
*Ddx3x* deletion in sensory neurons of adult mice causes spontaneous “mad itch”. **a,** Representative images of self-inflicted skin lesions on the face, flank and nape of *Ddx3x^BRN3A^*female mice, housed separately, 6 weeks after tamoxifen treatment. **b**, Schematics depicting the various behavioral events which contribute to the eventual skin lesions collectively referred to as “mad itch”. **c,d,** Time spent scratching, wiping, shaking, biting (**c**) and the number of such behavioral events (**d**) during a one-minute observational period in control *Ddx3x^WT^* (N=8) and littermate *Ddx3x^BRN3A^* (N=7) female mice 6 weeks after tamoxifen treatment. **e,** Top: prevalence of skin lesions in *Ddx3x^BRN3A^*female mice treated with tamoxifen or vehicle (oil). Bottom: schematic showing the time course of *Ddx3x^BRN3A^* female mice treated with tamoxifen and when the skin lesions typically appear. **f**,**g,** Representative immunofluorescence images of sensory neurons in DRG tissue from *Ddx3x^BRN3A^* female mice 6 weeks after treatment with tamoxifen or vehicle (oil) stained with anti-CGRP and IB4-lectin (**f**) and quantification of the number of each neuronal subtype (**g**). CGRP, calcitonin gene-related peptide; IB4, isolectin-B4. Scale bar, 100µm. Individual mice for each genotype are shown (c,d,g). Data are shown as means ± s.e.m. Two-tailed unpaired Student’s t test (c,d); Multiple t-test (g). *P < 0.05; ***P < 0.001; ****P < 0.0001; NS, not significant.

### DDX3X regulates mRNA metabolism and neurite architecture in sensory neurons

Our data so far showed that PRV infections result in a reduction of DDX3X levels and that loss of *Ddx3x* results in death of small-diameter, peripheral sensory neurons, suggesting that PRV infections might result in neuronal death via somal depletion of DDX3X. We first used RNAseq on DRGs isolated from *Ddx3x^BRN3A^* female mice treated with tamoxifen to investigate why the neurons eventually die. To assess the earliest changes caused by deletion of *Ddx3x*, we first performed a time-course to find the optimal period for tissue extraction when DDX3X protein levels were significantly reduced while neuronal integrity was preserved (**Figure S4a-c**). Using the *YFP^floxSTOPflox^* system confirmed that expression of *Brn3A*-*Cre* is specifically confined to peripheral DRG neurons and not spinal cord or brain (**Figure S4b**). RNAseq expression analysis validated that *Ddx3x* levels were significantly reduced in DRG tissue 2 days after the final tamoxifen treatment (**Figure S5a**).We and others have shown that *Ddx3x* plays essential roles in RNA metabolism and mRNA processing and is therefore vital for cell viability (Chen et al., 2016a; Szappanos et al., 2018; Tsherniak et al., 2017; Yu et al., 2016). Indeed, molecular functions associated with mRNA processing were the top dysregulated processes which were significantly downregulated upon *Ddx3x* deletion in sensory neurons **(Figure 3a; Figure S5b**).

**Figure 3.**
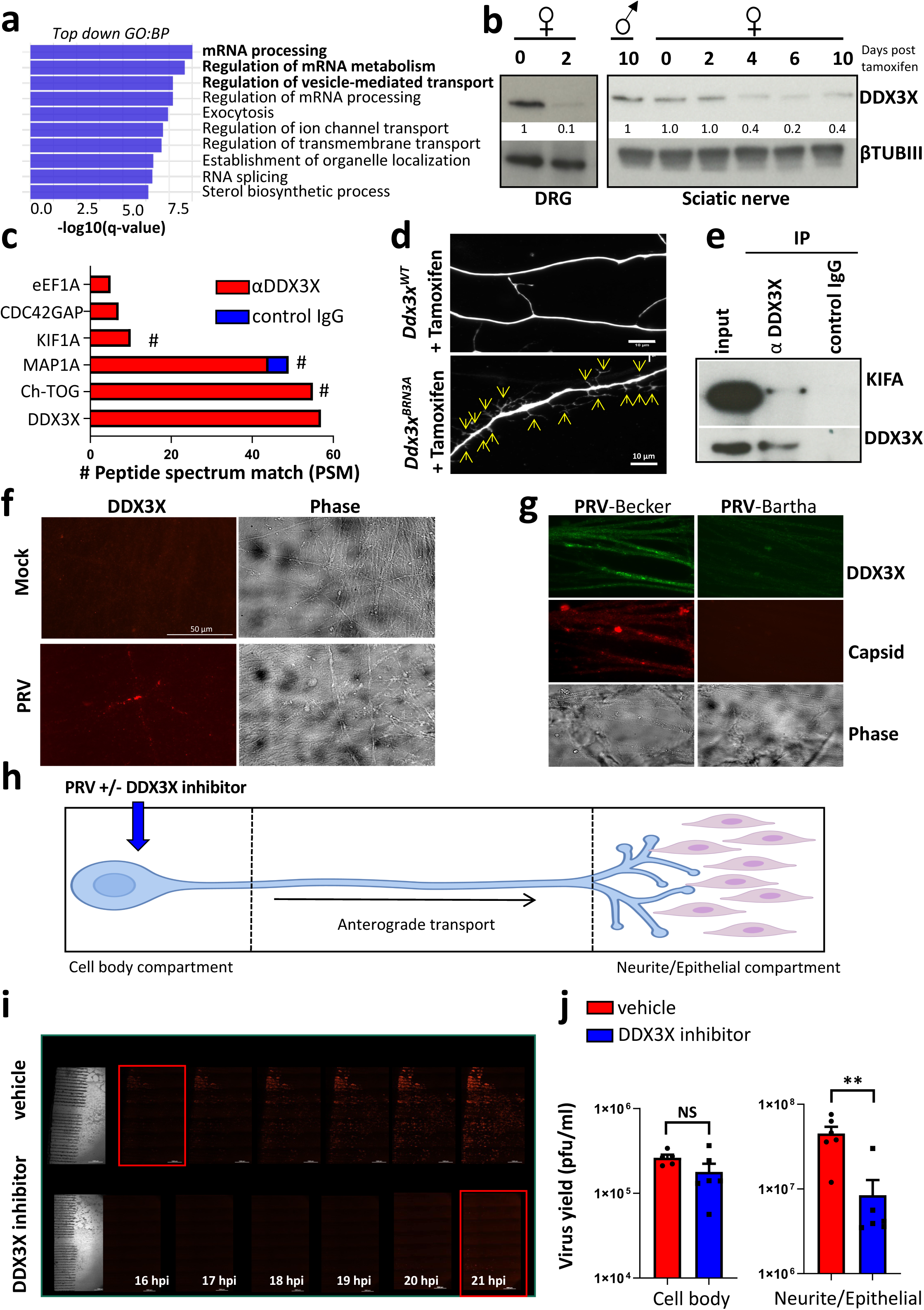
PRV hijacks DDX3X for axonal transport and spread. **a,** Table depicting the top 10 biological processes (BP) from gene ontogeny (GO) analysis of the downregulated transcripts in the sensory neuron specific *Ddx3x* knock-out mice comparing RNAseq data of DRG tissue from control *Ddx3x^WT^* and *Ddx3x^BRN3A^* female mice 6 weeks after treatment with tamoxifen. **b,** Western blot of DDX3X protein levels in DRG and sciatic nerve axonal tissue from *Ddx3x^BRN3A^* mice after treatment with tamoxifen for the indicated times. βTUBIII is used as a loading control. **c,** Proteins identified by mass spectrometry after immunoprecipitation of DRG tissue with anti-DDX3X and control IgG antibodies. Asterisks indicate proteins associated with axonal transport. **d,** Representative immunofluorescence of neurites from cultured DRG neurons from control *Ddx3x^WT^* and *Ddx3x^BRN3A^*female mice 3 weeks after treatment with tamoxifen. Arrows point to dendritic spines. Neurites were stained using βTUBIII. **e,** Western blot for KIF1A and DDX3X following immunoprecipitation of DRG tissue with anti-DDX3X and control IgG antibodies. IP, immunoprecipitation. **f,** Representative immunofluorescence images of neurites from mock- and PRV-infected SCG neurons 12 hours after infection showing DDX3X localization in the neurites. Phase images are also shown. **g,** Representative immunofluorescence images of DDX3X and labelled-PRV capsid in PRV-Becker-infected and PRV-Bartha-infected axons 12 hours post infection in SCG neurons. Phase images are also shown. **h,** Schematic depicting the tri-chamber system to investigate PRV anterograde transport to infect pig PK15 epithelial cells (neurite/epithelial compartment). PRV and the DDX3X inhibitor (called 16d) are added to the cell body compartment. **i,j,** Representative immunofluorescence images of labelled- PRV in pig PK15 epithelial cells after cell body infection with PRV in the tri-chamber system. Vehicle and DDX3X inhibitor were added one hour after PRV infection. Images were acquired at indicated times (hours) post infection (**i**). Quantification of PRV yield from the cell body and neurite/epithelial compartments 24 hours post infection after vehicle and DDX3X inhibitor treatments (**j**). Individual samples for each experiment are shown (j). Data are shown as means ± s.e.m. Two-tailed unpaired Student’s t test (j). **P < 0.01; NS, not significant.

Of note, there exists a homologue of *DDX3X* on the non-recombining region of the Y-chromosome, called *DDX3Y*, which is one of the very few genes maintained on the Y chromosome in evolution. While X-chromosome encoded DDX3X is ubiquitously expressed, DDX3Y protein expression was originally thought to be confined to the male germline (Ditton et al., 2004), but recent evidence suggests this not to be the case (Szappanos et al., 2018). Their high degree of similarity (over 90% similar at the amino acid level) supports the idea that DDX3X and DDX3Y are functionally redundant (Sekiguchi et al., 2004). Indeed, *Ddx3x^BRN3A^* male mice treated with tamoxifen did not display any signs of self-mutilation or “mad itch” behaviors. We further observed *Ddx3y* expression in peripheral sensory neurons of male mice, apparently compensating for the loss of *Ddx3x* (**Figure S5a**). The levels of other *Ddx* family members were unaltered upon *Ddx3x* deficiency (**Figure S5a**). These data provide *in vivo* organismal evidence that the Y-chromosome encoded DDX3Y homologue can functionally compensate for the loss of DDX3X.

When we analyzed gene sets which were perturbed specifically in *Ddx3x*-deficient sensory neurons (**Figure 3a; Figure S5b)**, axonal and vesicle transport were among the most significantly regulated biological processes, besides mRNA metabolism. This guided us to assess DDX3X localization within the neurons. Although the vast majority of DDX3X is localized to the soma, or cell bodies, of sensory neurons, it was also detected along neurites of cultured DRG neurons and present in the axons of sciatic nerves (**Figure 3b; Figure S6a**). We next performed co-immunoprecipitations with anti-DDX3X and control IgG antibodies from axonal sciatic nerve tissue followed by mass spectrometry (**Figure 3c**). The top candidates to bind DDX3X in the axons were proteins associated with the cytoskeleton such as ch-TOG (also known as cytoskeleton-associated protein 5) (Barr and Gergely, 2008) and microtubule associated protein (MAP)-1A (Fukuyama and Rapoport, 1995). Both proteins have been implicated in axonal and dendrite integrity in neurons (Halpain and Dehmelt, 2006; van der Vaart et al., 2012). Indeed, in the absence of DDX3X, the neurites of cultured DRGs displayed abnormal architecture with significantly increased dendritic spines (**Figure 3d**). Of note, DDX3X has been previously reported to play a role in RAC1-mediated dendritic spine homeostasis (Chen et al., 2016b; Zamboni et al., 2018). These data show that genetic inactivation of *Ddx3x* results in altered mRNA homeostasis and vesicle transport, affecting cell survival, and aboral architecture of sensory neurons.

### PRV hijacks DDX3X to facilitate its anterograde axonal transport

Another binding partner we identified for DDX3X in sciatic nerve axons was KIF1A, which was validated by co-immunoprecipitation (**Figure 3e**). KIF1A is a microtubule plus end-directed- motor kinesin-family protein involved in the anterograde transport of vesicles and organelles (Chiba et al., 2019; Okada et al., 1995). It has been shown that PRV virions use KIF1A to move anterogradely along axons to the synapse to facilitate spreading (Huang et al., 2020; Kramer et al., 2012). Importantly, in neuronal cultures, DDX3X robustly re-localized from the cell body to the neurites upon PRV infection (**Figure 3f; Figure S6b,c**). An attenuated strain of PRV (called PRV-Bartha) does not induce “mad itch” in infected animals compared to the wild type strain (called PRV-Becker). PRV Bartha lacks the US9-gE-gI proteins that are required for axonal sorting and efficient anterograde spread of virions along neuronal axons (Brideau et al., 2000; Husak et al., 2000; Yang et al., 1999). Indeed, PRV-Bartha infection of neurons did not lead to increased axonal localization of DDX3X (**Figure 3g),** further supporting the link between DDX3X and anterograde axonal transport of PRV virions.

To specifically examine whether DDX3X controls anterograde axonal transport of PRV, we used a compartmented tri-chamber neuronal culture system in which cell bodies (soma compartment) were fluidically and physically separated from neurites and porcine PK15 epithelial cells (neurite/epithelial compartment) (**Figure 3h**). Upon PRV infection of the cell bodies, the virus enters the neurons, replicates and progeny virion particles are transported anterogradely along the neurites to the synapses where they can then infect and replicate in the epithelial cells. To target DDX3X, we used a recently characterized small molecule DDX3X inhibitor called 16d (Brai et al., 2016). Cell bodies were infected with red-capsid labelled PRV (PRV180; mRFP-VP26). Vehicle or the DDX3X inhibitor were added 1-hour post infection to the cell body compartments. Red-capsid PRV first infected the epithelial cells ∼ 16 hours post infection and by 21 hours there was significant infection of the cells in the vehicle-treated samples (**Figure 3i**). By contrast, DDX3X inhibition significantly reduced the amount of PRV progeny that are anterogradely sorted and infecting the epithelial cells (**Figure 3i**). Viral yield measurements 24 hours post infection confirmed that DDX3X inhibition substantially reduced the amount of PRV reaching the epithelial cells (**Figure 3j; Supplementary Videos 3 and 4**). Viral titers and virion production from the cell body compartment were comparable between vehicle and DDX3X inhibition **(Fig. 3j; Supplementary Videos 5 and 6)**, indicating that inhibition of DDX3X does not significantly affect viral replication in the neuronal soma. It is important to note, that as opposed to genetic deletion of DDX3x, inhibition of DDX3X with 16d does not affect cellular viability on sensory neurons (Brai et al., 2016). HSV-1 is a human alphaherpesvirus which is also neurotropic, remains latent in the cell bodies of sensory neurons and upon reactivation travels along axons to keratinocytes where viral replication occurs leading to shedding and ‘cold sores’. Unlike PRV infection, HSV-1 infection of neurons does not induce robust axonal localization of DDX3X (**Figure S7a**). DDX3X has previously been linked to HSV-1 replication as well as host interferon response to HSV infection (Khadivjam et al., 2017; Soulat et al., 2008). Interestingly, we noticed that upon HSV-1 infection, DDX3X re- localizes from the perinuclear region to the nucleus (**Figure S7b**); something which is not seen upon PRV infection (**Figure S1a**). DDX3X inhibition had no apparent effect on the anterograde transport of HSV-1 (**Figure S7c,d**). These data demonstrate that PRV infections re-localize DDX3X from the sensory cell body to the axons to aid anterograde transport and subsequent spread of the virus.

### Inducible loss of adult small-diameter sensory neurons triggers “mad itch”

We next asked whether loss of the small-diameter DRG sensory neurons per se could account for the “mad itch” seen in tamoxifen-treated *Ddx3x^BRN3A^* female mice. To answer this, we used two genetically inducible systems to specifically ablate small-diameter DRG neurons based on the diphtheria toxin/receptor system (Buch et al., 2005; Ivanova et al., 2005). Floxed diphtheria toxin (DTx) transgenic (*DTx^floxSTOPflox^*) mice were bred to the *Brn3A-Cre-Ert* line for tamoxifen-inducible neuronal ablation (Buch et al., 2005) (**Figure 4a; Figure S8a**). Similarly, floxed diphtheria toxin receptor (DTR) transgenic mice (*DTR^floxSTOPflox^*) were bred to the *NaV1.8-Cre* line (generating *DTR^NaV1.8^*mice) for DTx-inducible neuronal ablation (Ivanova et al., 2005) (**Figure 4b,c; Figure S8b**). Both genetic strategies resulted in markedly impaired pain sensation (**Figure 4d,e**; **Figure S8c,d**), with accompanying 100% penetrance of skin lesions and self-mutilation in singly housed mice (**Figure 4f**; **Figure S8e-f**). Notably, the “mad itch” and extent of self-inflicted lesions were more apparent and intense in the *Ddx3x^BRN3A^* mice compared to the *DTR^NaV1.8^* model. Moreover, *DTR^NaV1.8^*but not *Ddx3x^BRN3A^* mice displayed significant weight loss and weakness after ablation of the neurons (**Figure S8h**). Importantly, neuronal ablation led to “mad itch” behaviors prior to lesion development (**Figure 4g**).

**Figure 4.**
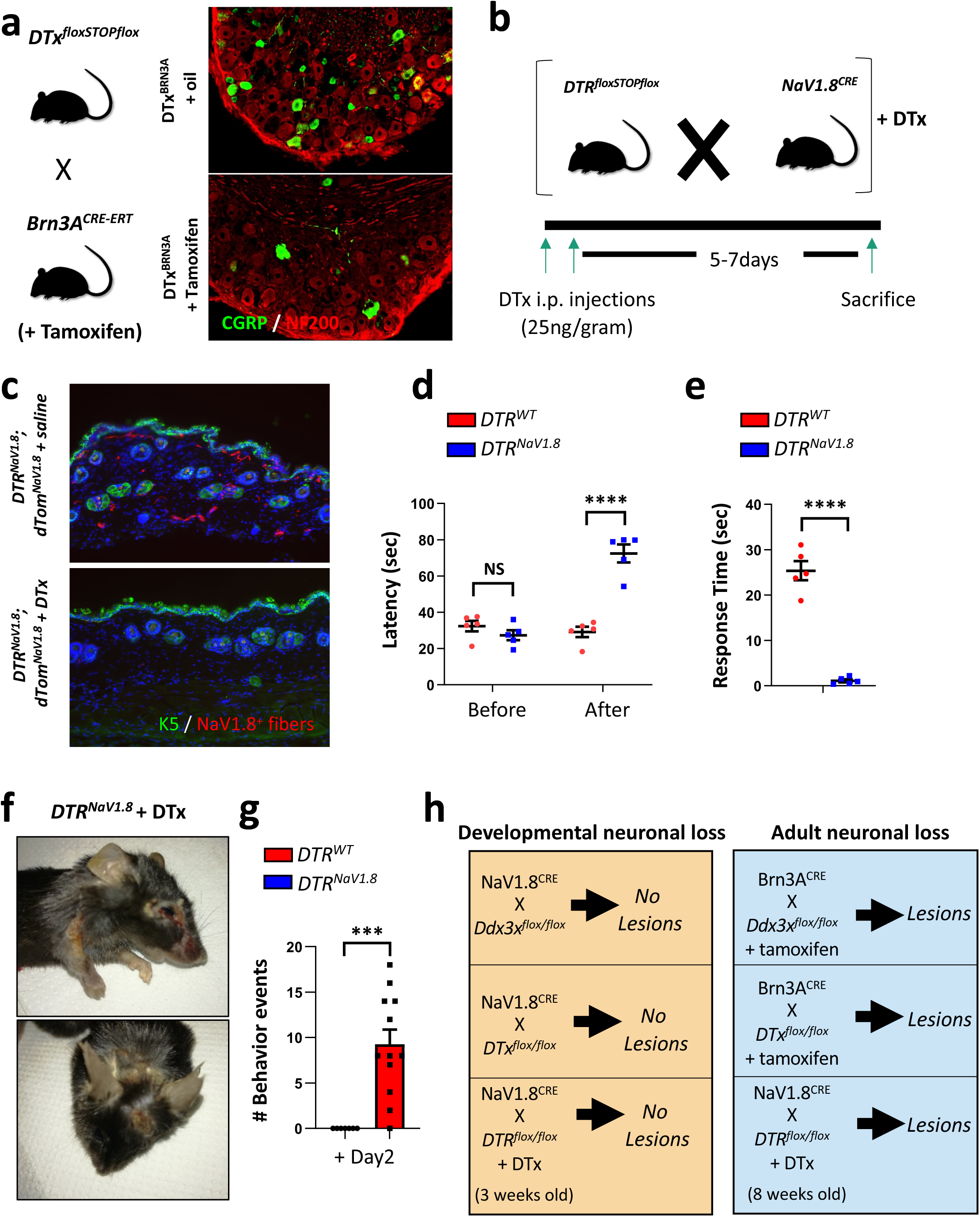
Loss of sensory neurons results in “mad itch”. **a,** Left: breeding scheme for inducible ablation of *Brn3a^+^*sensory neurons by inducible expression of diphtheria toxin (DTx) using tamoxifen treatment. Right: representative immunofluorescence images of CGRP^+^ (small diameter, unmyelinated sensory neurons) and NF200^+^ (large diameter, myelinated sensory neurons) neurons in DRG tissue from *DTx^BRN3A^* mice one week after treatment with tamoxifen or vehicle (oil). **b,** Breeding scheme to temporally ablate *NaV1.8^+^*sensory neurons by inducing diphtheria toxin receptor (DTR) followed by DTx treatment. **c,** Representative immunofluorescence images of skin tissue labelled with K14 (green), DAPI (blue) and the endogenous *NaV1.8-TdTomato* reporter (red) from *DTR^NaV1.8^,TdTomato^NaV1.8^*mice one week after treatment with DTx or vehicle (saline). **d**, Contact heat withdrawal latencies using the hotplate assay at 50°C for control *DTR^WT^* (N=5) and littermate *DTR^NaV1.8^* (N=5) mice before and five days after DTx treatment. The experiment was repeated an additional time with similar results. **e**, Nocifensive responses in a 3-minute time frame after intraplantar injection of 2 μg of capsaicin into the left hind paw of control *DTR^WT^* (N=5) and littermate *DTR^NaV1.8^*(N=5) mice five days after DTx treatment. **f,** Representative images of skin lesions on the face, nape and flank of singly-housed *DTR^NaV1.8^*mice 1 week after DTx treatment. **g,** Number of “mad itch” behavioral events (wiping, shaking, biting) during a two-minute observational period in control *DTR^WT^* (N=7) and littermate *DTR^NaV1.8^* (N=12) mice two days after DTx treatment. **h,** Summary of the prevalence of lesions and itch-like pathological behavior using various models in which sensory neurons were ablated early during development or temporally in the young (3 weeks old) and adult (all inducible mutants were >8 weeks of age at the start of treatment) mice. Individual mice for each genotype are shown (d,e,g). Data are shown as means ± s.e.m. Multiple t-test (d); two-tailed unpaired Student’s t test (e,g). ***P < 0.001; ****P < 0.0001; NS, not significant.

In both *Cre*-expressing lines, loss of small-diameter sensory neurons resulted in “mad itch” and self-mutilation, which intriguingly depended on the age of the mice when ablation occurred – younger mice failed to show any overt signs of itch and self-mutilation (**Figure 4h; Figure S8g**). This time-dependency of neuronal ablation was also confirmed by crossing *Ddx3x^flox/flox^*mice to the *NaV1.8-Cre* line which deletes *Ddx3x* and thereby ablates the small-diameter sensory neurons early in development (Ivanova et al., 2005) (**Figure S9a**). Similar to *Ddx3x^BRN3A^* female mice, *Ddx3x^NaV1.8^*female mice showed insensitivity to pain and, although a progressive loss of sensory neurons was observed by 4 weeks of age, no indication of “mad itch” was observed (**Figure 4h**; **Figure S9b-g**), indicating that neuronal loss prior to a certain age does not lead to the “mad itch”. Altogether these data show that loss of small-diameter sensory neurons in adulthood results in “mad itch”.

### No apparent role of immune cells in initiating the “mad itch”

One of the most common types of chronic itch in patients is atopic dermatitis (Langan et al., 2020). In this type of itch, the immune system and various immune-related cytokines play a major role in its pathogenesis (Langan et al., 2020). To assess whether the immune system initiates or propagates “mad itch”, we used our *DTR^NaV1.8^* model due to its rapid onset of self-inflicted skin lesions and ease to combine with additional genetic and pharmacological mouse lines (**Figure S10a**). Histological examination demonstrated strong immune cell infiltration in skin lesions after DTx treatment (**Figure S10b,c**). Moreover, systemic inflammation was also evident from enlarged spleens in *DTR^NaV1.8^*mice compared to *DTR^WT^* mice seven days after DTx treatment (**Figure S10d**). To examine whether an immune response precedes and initiates the manifestation of skin lesions, we analyzed various inflammatory markers in the serum of *DTR^WT^* and *DTR^NaV1.8^* mice 2 days after DTx treatment, i.e. before the appearance of any physical signs of self-inflicted skin lesions. Of the 15 selected pro-inflammatory markers, only IL-6 and G-CSF were significantly increased in the serum of *DTR^NaV1.8^*mice (**Figure 5a**). These data were confirmed in inducible *Ddx3x^BRN3A^* female mice in which serum IL-6 and G-CSF levels rapidly increased after tamoxifen treatment, again prior to any physical signs of self-mutilation (**Figure 5b,c**). Interestingly, IL-6, and to a lesser extent G-CSF, are both induced and released from injured DRG sensory neuronal cultures (**Figure 5d**). Serum IL-6 and G-CSF were also reported to be specifically increased upon PRV infection in mice (Laval et al., 2018). Moreover, we demonstrate that *Ddx3x* deletion also induces a robust injury response in DRG tissue as evidenced by increased levels of the prototypical early injury response protein Activating transcription factor 3, (ATF3) (**Figure 5e**) (Tsujino et al., 2000). This led us to investigate whether ATF3 is also increased in DRG neurons upon PRV infection. Indeed, nuclear ATF3 is substantially increased preferentially in small-diameter peripheral sensory neurons of the L4/L5 DRG tissue after hind-paw PRV infection in mice (**Figure S10e**), again highlighting the striking similarities between PRV infection and DRG small fiber-specific *Ddx3x* deletion.

**Figure 5.**
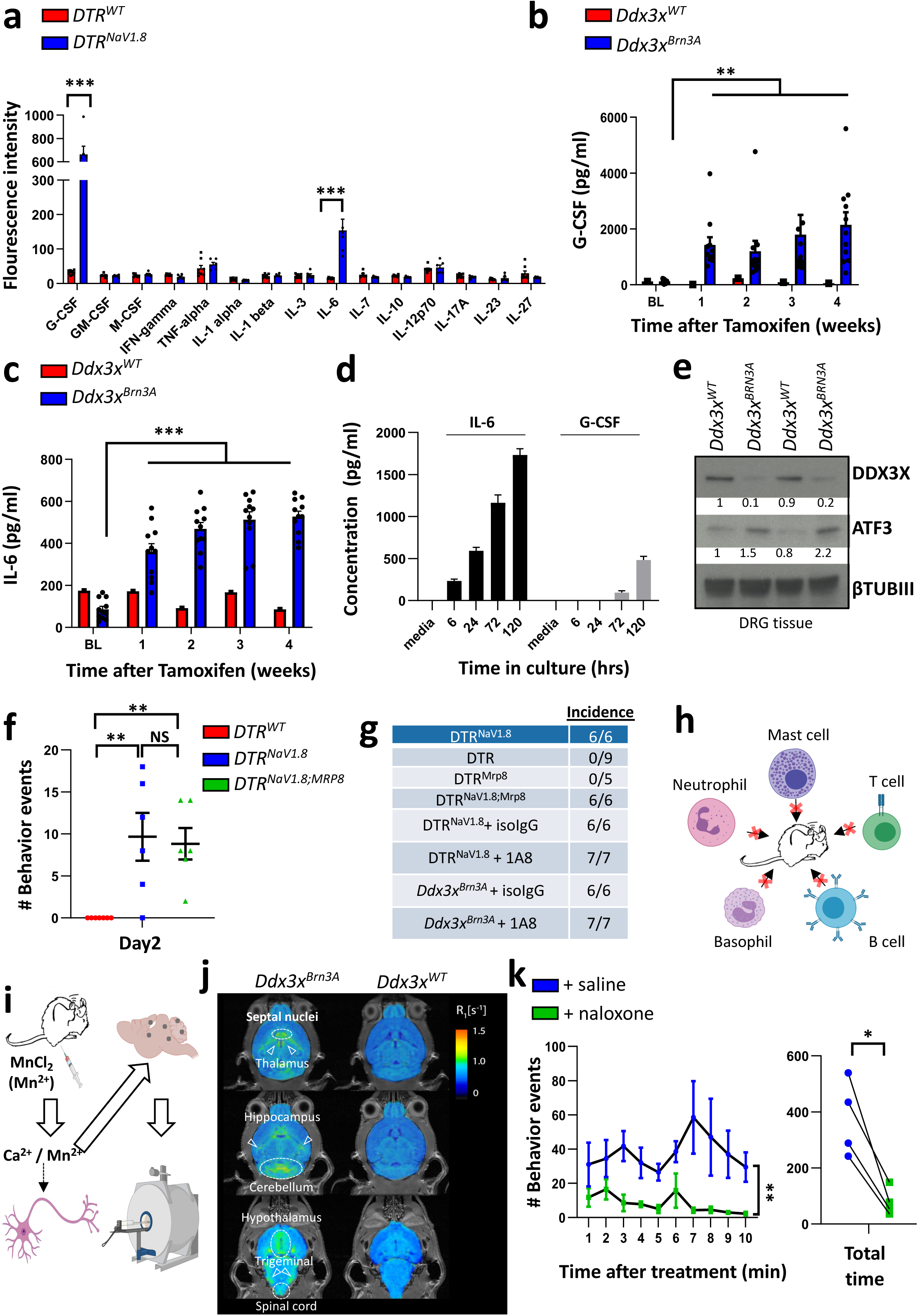
Addictive “mad itch”. **a,** Serum profile analysis of inflammatory cytokines from *DTR^WT^* (N = 8) and *DTR^NaV1.8^* (N = 6) mice after 2 days of DTx treatment. **b,c,** Quantification of G-CSF (**b**) and IL-6 (**c**) over time from the serum of *Ddx3x^WT^* (N=1) and littermate *Ddx3x^BRN3A^*(N=11) female mice before and after tamoxifen treatment. **d**, IL-6 and G-CSF levels in supernatant over time from wild type DRG neuronal cultures. **e,** Western blot analysis of DDX3X and ATF3 from DRG tissue of *Ddx3x^WT^*(N=2) and littermate *Ddx3x^BRN3A^* (N=2) female mice five days after tamoxifen treatment. βTUBIII is used as a loading control. **f,** Number of “mad itch” behavioral events (wiping, shaking, biting) during a two-minute observational period in control *DTR^WT^* (N=7), *DTR^NaV1.8^* (N=6) and *DTR^NaV1.8;MRP8^* (N=7) mice two days after DTx treatment. **g**, Table depicting the contribution of neutrophils to the incidence of skin lesions. **h,** Summary schematic showing the various immune cell subtypes which do not appear contribute to the prevalence of “mad itch” through genetic ablation and depletion studies. **i**, Schematic showing how manganese (Mn^2+^)-enhanced magnetic resonance imaging (MEMRI) is used to determine areas of neuronal activation using the paramagnetic Mn^2+^ as an analog for calcium uptake in excitable neurons. **j,** Representative R1-maps for *Ddx3x^BRN3A^*and *Ddx3x^WT^* female mice 24h following MnCl^2^ injections. Maps were pseudocolored based on R1 values (see color scale) and overlaid over anatomical scans. In *Ddx3x^BRN3A^* female mice there was ∼30% increase in R1 values in septal nuclei, thalamus, hippocampus, cerebellum, trigeminal nuclei and spinal cord. *Ddx3x^WT^* mice did not show similar increases in R1 values, thus the highlighted areas represent neuronal activity in response to “mad itch”. **k,** Left: time course of the effect of saline and naloxone (20mg/kg) on the number of behavioral events associated with “mad itch” in a 10- minute observational period in *Ddx3x^BRN3A^* female mice 6 weeks after treatment with tamoxifen. Each mouse was analysed on two consecutive days at the same time of day with saline injected on the first and naloxone on the second. Right: total number of behavioral events recorded in a 10-minute period. Individual mice for each genotype are shown (a,b,c,f,k). Data are shown as means ± s.e.m. Multiple t-test (a); Two-way ANOVA with a Dunnett’s multiple comparison test to baseline (b,c); One-way ANOVA with Tukey’s multiple comparison test (f); Two-way ANOVA (k, left); Two-tailed paired Student’s t test (k, right). *P < 0.05; **P < 0.01; ***P < 0.001; NS, not significant.

IL-6 and G-CSF both play a key role in neutrophil recruitment (Yan et al., 2013; Zhang et al., 1998). To investigate whether neutrophils play a role in “mad itch” development in our mouse model, we genetically ablated neutrophils at the same time as the sensory neurons, using the neutrophil-specific *MRP8-Cre* deleter line (Passegué et al., 2004) (**Figure S11a**). As expected, DTx treatment resulted in a dramatic loss of neutrophils (**Figure S11b**). A second approach employed to deplete neutrophils was to use the anti-Ly6G depleting antibody 1A8 in *DTR^NaV1.8^* and inducible *Ddx3x^BRN3A^* female mice (Brown et al., 2004) which also significantly reduced neutrophil numbers (**Figure S11c**). However, neither approach affected the incidence of “mad itch”-like behaviors, self-mutilation or skin lesions in *DTR^NaV1.8^* mice compared to vehicle-treatment (**Figure 5f,g**). We next investigated whether the adaptive immune system, namely T and B cells, initiate the “mad itch”. Again, we used two different approaches to selectively ablate T and B cells; breeding the *DTR^NaV1.8^* line to the *Rag1-expressing Cre* line (McCormack et al., 2003) and, secondly, to cross the *DTR^NaV1.8^* line onto a Rag2^KO^ background (Hao and Rajewsky, 2001) – both strategies effectively ablate T and B cells (**Figure S11d-f**). Although spleen weights were reduced in *DTR^NaV1.8; Rag1^*and *DTR^NaV1.8^;Rag2^KO^* mice compared to their respective littermates 7 days after DTx treatment, there was no effect on the incidence of skin lesions or “mad itch” (**Figure S11g-j**).

One of the most potent and classic mediators for pathological itch is histamine (Buddenkotte et al., 2010). IgE binds to the FcεRI receptor on mast cells and basophils to induce degranulation and release of histamine (Borriello et al., 2017). Both serum histamine and IgE levels were significantly upregulated in *DTR^NaV1.8^*mice 7 days after DTx treatment (**Figure S11k**). To investigate the contribution of mast cells and basophils to “mad itch” induction, we genetically ablated these cell types using the *Cpa3-Cre* expressing line (Lilla et al., 2011) (**Figure S11l,m**). However, removal of mast cells and basophils had no effect on skin lesion incidence in our *DTR^NaV1.8^* model (**Figure S11n**). Based on these data (**Figure 5h**), the role of the immune system appears to be secondary to the self-mutilation and ensuing tissue damage, though we cannot exclude redundancies among distinct immune cells.

### Self-mutilation is driven by an addiction to the pleasure of “mad itch”

Since the immune effects are likely secondary to the self-inflicted tissue damage, we wanted to investigate the brain circuits of mice undergoing “mad itch”. To accomplish this, we used manganese-enhanced magnetic resonance imaging (MEMRI) (Lin and Koretsky, 1997) (**Figure 5i**). This involved injecting a manganese (Mn^2+^) chloride solution into *Ddx3x^WT^* and *Ddx3x^BRN3A^* mice 6 weeks after tamoxifen treatment when itch-related behaviors and self-mutilation were evident. Manganese ions are paramagnetic and conduct through calcium-permeable channels on activated neurons (Lin and Koretsky, 1997). Mn^2+^ is a T1-contrast agent that upon accumulation in the neurons leads to reduction in longitudinal relaxation time (T1) and an increase in relativity rate R1 (R1=1/T1). Using a 15.2 Tesla MRI, MEMRI allowed us to image stimulated brain regions irrespective of changes in blood flow and thereby identify areas activated during a 24-hour timeline analysis of “mad itch”.

Several major brain regions were markedly different between free-moving *Ddx3x^WT^* and *Ddx3x^BRN3A^* mice during the 24-hour period (**Figure 5j**). In *Ddx3x^BRN3A^*mice, expectedly (Chen and Sun, 2020; Dong and Dong, 2018; Lipshetz et al., 2018), areas associated with sensory processing were activated – namely the spinal cord, trigeminal nerves and thalamus (**Figure 5j**). Moreover, increased activation of the hippocampus, hypothalamus and cerebellum were noted, regions previously associated with neuropathic itch in patients (Ju and Yosipovitch, 2020). Interestingly, in addition to these prototypic circuits of itch sensation, we also detected strong activation of the septal nuclei in *Ddx3x^BRN3A^* mice undergoing “mad itch” (**Figure 5j**). The septal nuclei are an area in the brain associated with the experience of pleasure, reward and reinforcement to sex or food, as well as addiction in rodents and humans (Berridge and Kringelbach, 2015), (Olds and Milner, 1954),(HEATH, 1963; Pantazis and Aston-Jones, 2020). The idea of the *Ddx3x^BRN3A^* mice being ‘addicted’ to the itch along with the well-documented clinical manifestation of opioid-induced itch seen in patients and recreational drug users (Bergasa et al., 1995; Reich and Szepietowski, 2010), prompted up to test whether opioid signaling is involved in the “mad itch”. Naloxone, a non-selective, competitive opioid receptor antagonist, dramatically reduced the amount of itch-related behavioral events in *Ddx3x^BRN3A^* mice (**Figure 5k; Figure S12**). Our data indicate that “mad itch” triggers the pleasure and reward center in the brain which might explain the intense self-inflicted deep tissue damage and skin lesions observed in the *Ddx3x^BRN3A^*mice. Inhibition of opioid receptors significantly diminishes the “mad itch” and self-mutilating behavior, providing a druggable intervention.

### Loss of sensory neurons leads to thymic atrophy and reduced T cell numbers

Although the immune system does not seem to play a major role in the initiation of the observed “mad itch”, viruses frequently divert host immunity to facilitate replication and viral spreading (Alcami and Koszinowski, 2000). We therefore asked whether ablation of sensory neurons per se might contribute to immune alterations. Since “mad itch” was associated with enlarged spleens, in part dependent on the presence of T cells, we first characterized immune cells in the spleen. Similar to *DTR^NaV1.8^* mice (**Figure S10d**), the spleens of *Ddx3x^BRN3A^* female mice were also increased in size compared to littermate *Ddx3x^WT^* control mice 6 weeks after tamoxifen administration (**Figure S13a**). This increase in spleen size was largely attributable to increased numbers of CD11b^+^ monocytes and neutrophils (**Figure S13b**). Despite the larger size, the spleens of *Ddx3x^BRN3A^*female mice 6 weeks after tamoxifen treatment contained significantly fewer CD4^+^ and CD8^+^ T cells, while B cell numbers remained largely unaffected (**Figure S13c**). We next assessed spleens from *Ddx3x^BRN3A^* female mice 4 weeks after tamoxifen treatment, a time point prior to the development of the skin lesions. However, at this time point we failed to observe any significant differences compared to control *Ddx3x^WT^* mice, although reduced numbers of sensory neurons were already apparent (**Figure S13d-f**).

At this time point, our MRI data also showed a strong activation of the hypothalamus in the ‘mad itch’ *Ddx3x^BRN3A^*mice (**Figure 5j**). The hypothalamus is a command center for the stress response and part of the HPA (hypothalamus-pituitary-adrenocortical) axis which controls secretion and levels of the stress hormone corticosterone in mice. Indeed, we observed a substantial increase in serum corticosterone levels in tamoxifen-treated *Ddx3x^BRN3A^* female mice (**Figure 6a**). Corticosterone has been demonstrated to induce thymic atrophy and death of thymic immune cells (Savino, 2006); we observed a marked thymic atrophy and altered T cell development affecting double-positive (DP) CD4^+^CD8^+^ and single positive (SP) CD4^+^ or CD8^+^ thymocytes in the *Ddx3x^BRN3A^*female mice 4 weeks after tamoxifen treatment (**Figure 6b,c**), prior to skin lesions and any apparent changes in the spleen. We also observed extensive thymic atrophy and markedly impaired T cell development in *Ddx3x^BRN3A^* female mice 6 weeks after tamoxifen treatment (**Figure S13g-j**). Progressive thymic atrophy was also detected in *DTR^NaV1.8^* mice treated with DTx (**Figure 6d**), accompanied by a marked reduction in developing thymocytes, particularly in DP CD4^+^CD8^+^ cells, before any skin lesions were apparent (**Figure 6e-h; Figure S14a**). The sensory neuronal ablation also resulted in progressive alterations in thymic architecture and increased levels of serum corticosterone (**Figure S14b,c**).

**Figure 6.**
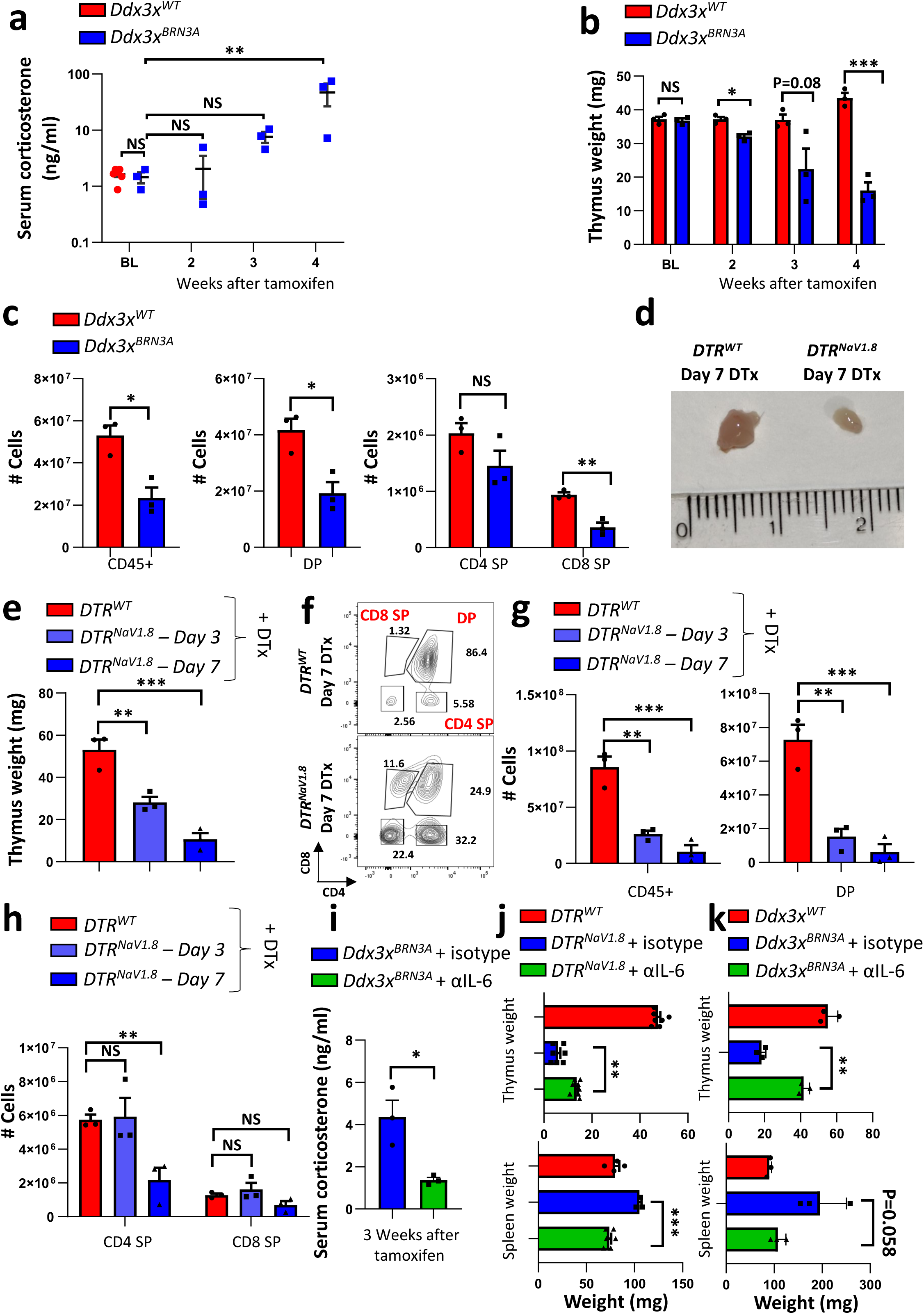
Sensory neuronal loss leads to thymic atrophy and systemic reduction in T cells. **a,** Serum corticosterone levels in control *Ddx3x^WT^* (N=6) mice at baseline and in littermate *Ddx3x^BRN3A^* (N=3) female mice at baseline and each week after tamoxifen treatment. **b,** Thymus weights of control *Ddx3x^WT^* (N=3) and littermate *Ddx3x^BRN3A^* (N=3) female mice at baseline and the indicated time points after tamoxifen treatment. **c,** Cell number of all thymocytes (CD45^+^) and the indicated thymic subpopulations of control *Ddx3x^WT^* (N=3) and littermate *Ddx3x^BRN3A^*(N=3) female mice at 4 weeks after tamoxifen treatment. **d,** Representative image of the thymus from 8-week-old control *DTR^WT^* and littermate *DTR^NaV1.8^*mice 7 days after DTx treatment. **e,** Thymus weights from 8-week-old control *DTR^WT^* (N=3) and littermate *DTR^NaV1.8^* mice analysed 3 (N=3) and 7 (N=3) days after DTx treatment. **f,** Representative FACS blots of CD4 and CD8 staining in the thymus of 8-week-old control *DTR^WT^* and littermate *DTR^NaV1.8^*mice 7 days after DTx treatment. SP, single positive; DP, double positive. **g,h,** Numbers of the various indicated cell types in the thymus of 8-week-old control *DTR^WT^* (N=3) and littermate *DTR^NaV1.8^* mice assayed 3 (N=3) and 7 (N=3) days after DTx treatment. **i**,Serum corticosterone levels in tamoxifen-treated *Ddx3x^BRN3A^*mice treated with isotype control antibody (N=3) and *Ddx3x^BRN3A^* treated with 100μg (for 5 consecutive days one week post tamoxifen). Serum levels were collected three weeks after tamoxifen treatment. **j,** Thymus and spleen weights of control *DTR^WT^* (N=7), *DTR^NaV1.8^* treated with isotype control antibody (N=7) and *DTR^NaV1.8^* treated with 100μg (every day starting one day after first DTx administration) anti-IL-6 antibody (N=9) 7 days after DTx treatment. **k**, Thymus and spleen weights of control *Ddx3x^WT^* (N=3), *Ddx3x^BRN3A^* treated with isotype control antibody (N=3) and *Ddx3x^BRN3A^* treated with 100μg (for 5 consecutive days one week post tamoxifen) anti-IL-6 antibody (N=3) 7 weeks after tamoxifen treatment. Individual mice for each genotype are shown (a,b,c,e,g,h,i,j,k). Data are shown as means ± s.e.m. Two-way ANOVA with Dunnett’s multiple comparison test (a). Two-tailed unpaired Student’s t test (c,j,i,k); Multiple t-test (b,c); One-way ANOVA with a Dunnett’s multiple comparison test (e,g,h); *P < 0.05; **P < 0.01; ***P < 0.001; NS, not significant.

Cytokines, in particular IL-6, have been previously shown to be involved in thymic atrophy (Ansari and Liu, 2017; Carbajosa et al., 2017; Sempowski et al., 2000). Moreover, IL-6 has been shown to activate the HPA axis and lead to increased corticosterone levels (KAPCALA et al., 1995; Späth-Schwalbe et al., 1994). Since increased serum levels of IL-6 in our *DTR^NaV1.8^* and *Ddx3x^BRN3A^* models precede the increase in corticosterone (**Figure 5c; Figure 6a**), we investigated whether this cytokine might be also mechanistically involved in thymic atrophy following neuronal loss. A five-day anti-IL-6 treatment regimen administered 1 week after tamoxifen injection, substantially reduced serum corticosterone levels after two additional weeks compared to isotype treatment (**Figure 6i**). Importantly, anti-IL-6 antibody treatment partially mitigated the reduction in thymic and spleen weights in both *DTR^NaV1.8^* mice treated with DTx (**Figure 6j**) and *Ddx3x^BRN3A^* female mice after tamoxifen treatment (**Figure 6k**). Furthermore, anti-IL-6 antibody treatment in part rescued the reduction in thymocyte populations as well as the decrease of peripheral T cells in the spleen using both models (**Figure S14d-f**), suggesting that the increased serum IL-6 plays a role in sensory neuron ablation-mediated thymic atrophy, though other mechanisms must be also operational. Anti-IL-6 antibody treatment did not however affect the “mad itch” dependent skin lesion development (**Figure S14g**). Altogether, these data demonstrate that by reducing DDX3X levels in sensory neurons, the ensuing neuronal loss triggers activation of the HPA axis, possibly via IL-6, which causes corticosterone-induced thymic atrophy leading to diminished numbers of developing thymocytes and peripheral CD4^+^ and CD8^+^ T cells.

## Discussion

Our results provide a framework (**Figure S15a,b**) for how PRV, through manipulation of DDX3X function, localization and stability, induces a range of neurological and immunological effects to subvert and co-opt host responses to ultimately aid the spread of the virus though itch- induced skin lesions.

Itch is a highly conserved physiological response - mammals, birds, zebrafish and even fruit flies have displayed site-directed, itch-like behaviors (Esancy et al., 2018; Li et al., 2016). Acute itch is caused by chemical or mechanical stimuli such as mosquito bites which can be effectively treated by anti-histamines. Under certain pathophysiological conditions, such as kidney dysfunctions or particular cancers, itch can become chronic. Neuropathic itch, which represents a subtype (approximately 8%) of chronic itch, is caused by an intrinsic malfunction of neuronal circuitry (Stumpf and Ständer, 2013). Alphaherpesvirus infections, such as VZV, are one of the main causes for severe neuropathic itch seen in humans (Steinhoff et al., 2018). The neurotrophic alphaherpesvirus PRV inflicts a “mad itch” in animals and has been used to study the effects of herpes virus-induced infection and neuropathic itch. PRV almost exclusively infects non-human mammals though there are some rare case reports to suggest human infection may occur (Guo et al., 2021; Wang et al., 2020). Here we show that upon infection of neurons, PRV induces the re-localization of DDX3X from the neuronal cell body to the axons and ultimately the reduction of DDX3X levels. In axons, DDX3X plays a role in the anterograde transport of PRV and the virus requires DDX3X to spread from the cell body to the synapse to facilitate its spread. PRV has three envelope proteins (gE-gI-Us9) that organize the lipid rafts so that mature virions can recruit KIF1A to sort into axons to allow efficient anterograde axonal transport and spread (Brideau et al., 2000; Husak et al., 2000). We now show that DDX3X binds KIF1A, as well as other mediators of axonal transport and integrity, suggesting that PRV hijacks this network to enable its anterograde axonal spread. Unlike PRV, DDX3X inhibition did not affect the anterograde transport of the related human herpes virus, HSV-1, a herpesvirus which is not known for causing neuropathic itch. Although PRV and HSV-1 share similar sorting strategies in the nervous system, there has been differences reported by several groups: while PRV Us9-KIF1A (kinesin 3) interaction seems to be the major player in anterograde sorting, in HSV-1 conventional kinesin (kinesin 1) interaction with Us9-gE-gI complex has been reported to be essential. Moreover, there have been observations of separate anterograde transport of HSV-1 sub-virion structures into axons, while the “married model” of sorting, where virion assembly is completed in the cell body before sorting, is mostly observed for PRV (DuRaine and Johnson, 2021; Kratchmarov et al., 2012). Moreover, DDX3X did not noticeably re-localize to the axons upon HSV-1 infection further arguing against its role in aiding anterograde transport for HSV-1. This differential response of DDX3X re-localization to HSV and PRV infections may explain why neurons are more resistant to HSV than PRV and why PRV infections can result in self-mutilating “mad itch”. The DDX3X inhibitor used in this study, 16d, specifically blocks the RNA binding domain of DDX3X (Brai et al., 2019). Importantly, the 16d-mediated inhibition is not lethal to cells or animals in vivo (Brai et al., 2019) and its inhibition specifically blocks PRV anterograde transport and not HSV. How inhibiting the RNA binding domain affects PRV transport or interaction with KIF1A needs further investigating.

PRV infections cause death of peripheral sensory neurons (Laval and Enquist, 2020). We modeled PRV-induced DDX3X dysregulation by specifically and temporally ablating *Ddx3x* in adult mouse small-diameter sensory neurons to mimic PRV infection. Inducible loss of *Ddx3x* indeed led to progressive loss of sensory neurons. In our *DDx3x^BRN3A^*model, the loss of small-fiber sensory neurons was accompanied by loss of pain sensation which by extension might aid in the self-mutilation. Moreover, our genetic models of *Ddx3x* deletion paralleled several key characteristics observed in PRV-induced “mad itch” in multiple animal species (**Table S1**). Neuropathic itch is typically associated with a decrease in the density of small-diameter nerve fibers that innervate the epidermis (Misery et al., 2014; Pereira et al., 2016). For example, in VZV-induced postherpetic itch (PHI), only 5% of epidermal sensory nerve fibers remained (Oaklander, 2011; Oaklander et al., 2002; Steinhoff et al., 2018). In a study of over a hundred adults with shingles, 17% developed PHI (Oaklander et al., 2003), with varying degrees of itch intensity; some described as a “compulsive scratching” of the affected areas leading to severe deep tissue damage (Oaklander et al., 2002; Patnaik et al., 2021). VZV only infects humans but its numerous similarities to PRV in causing neuropathic itch may hint at a role for DDX3X, though this needs to be investigated.

Moreover, a recent case study of neuropathic itch may have resulted from an injury to the DRG tissue during a DRG stimulator trial for chronic pain (Strand et al., 2021). Using two genetic approaches to inducibly ablate sensory neurons, we confirmed that small-fiber peripheral neurodegeneration causes intense itch-like behaviors and self-mutilation. Interestingly, there is a temporal element to itch induction from loss of neurons; when *Ddx3x* is ablated during development (*Ddx3x^flox^* X NaV1.8-Cre), or when sensory neurons are ablated during development (DTx^flox^ X NaV1.8-Cre mice) or when DTx is administered to 2-week-old *DTR^NaV1.8-Cre^* mice, there is no accompanying itch. This also rules out the possibility that loss of pain combined with self-grooming, could lead to self-inflicted skin lesions. An initiating signal from injured sensory neurons in the adult is required for “mad itch” development. Why this itch behavior is age dependent in our model requires additional experiments. A recent report modeling a sodium channel NaV_1.7_ gain-of-function mutation demonstrated unexplained self-inflicted skin lesions in older mice (Wimalasena NK, 2023). The authors describe an age-dependent nociceptive insensitivity with a specific loss of tyrosine hydroxylase-positive (TH^+^) c-fiber low threshold mechanoreceptors (C-LTMRs). We posit that this loss of small-diameter TH^+^ neurons may be the responsible subtype for the initiation of the “mad itch”.

Chronic itch conditions are strongly linked to dysregulated immune cell activation with neutrophils, mast cells, basophils, T and B cells all playing prominent roles in initiating and propagating ‘itch’ signals via sensory neurons (Mack and Kim, 2018). Using a combination of genetic ablation and antibody-mediated depletion strategies, we have effectively ruled out a role of these immune cells in the observed “mad itch” skin lesions induced by sensory neuronal loss, suggesting the “mad itch” we observe is driven neuronally. Manganese enhanced magnetic resonance imaging allowed us to determine what brain regions are activated in *DDx3x^BRN3A^* mice during a 24-hour “mad itch” observational window. Interestingly, in addition to areas previously associated with itch sensation such as the spinal cord, trigeminal nerves, hippocampus and thalamus (Chen and Sun, 2020; Dong and Dong, 2018; Lipshetz et al., 2018), the septal nuclei were hyperactivated in our “mad itch” mice. This region critically regulates pleasure, addiction, and reinforcement behavior, suggesting that addiction-pleasure might be involved in the observed itch-related behaviors. Indeed, according to the 16^th^ century philosopher *Michel de Montaigne,* ‘scratching is one of nature’s sweetest gratifications’, and mirrors the description of PRV-infected animals where the itch “escalated in severity until the animals directed the entirety of their energy to scratching and biting” the affected areas (Brittle et al., 2004). Moreover, itch is a common side effect of opioids used in the treatment of intractable or post-operative pain (Ko, 2015; Yamamoto and Sugimoto, 2010). Generalized pruritus is also a well-known symptom of patients with chronic opioid abuse. While opioids can cause histamine release, these patients are commonly resistant to anti-histamines, strongly suggesting a direct mediation by opioids (Lipman and Yosipovitch, 2021). Importantly, naloxone treatment significantly reduced the “mad itch” in *DDx3x^BRN3A^* mice showing that the itch is driven by opioid receptor signaling. Our data imply a fundamental link between “mad itch” and activation of brain pleasure/addiction centers to maintain “mad itch” behaviors through endogenous opioid signals.

Pathogens, in particular certain viruses, have been shown to induce acute thymic atrophy as a strategy to divert host defenses during infection (Albano et al., 2019; Ansari and Liu, 2017). Importantly, herpes viruses have been shown to induce thymic atrophy in chicken (Gimeno et al., 2011), cattle (Romero-Palomo et al., 2015), seals (Gulland et al., 1997) and also in a case study in a human neonate (Watanabe et al., 1984). PRV infection has also previously been linked to thymic atrophy (Narita and Ishi, 2006). Recently, sensory neuro-innervation of the peripheral lymphoid organs has been demonstrated (Hu et al., 2020; Huang et al., 2021); however, such innervation was not observed in the thymus suggesting indirect effects from sensory neuron loss may affect thymic architecture (Gautron et al., 2011). We demonstrate that IL-6 and G-CSF are massively increased in the serum after *Ddx3x* deletion in sensory neurons as well as after ablation of sensory neurons, phenocopying the increased levels of these two cytokines after PRV infection in mice (Laval et al., 2018). Both cytokines, but in particular IL-6, are increased in injured DRG tissue (Dubový et al., 2013; Ogawa et al., 2014) and in cultured DRG neurons. Loss of *Ddx3x* also induced a rapid injury response evident by increased ATF3 levels in DRG tissue early after tamoxifen treatment and we also show that this effect is mirrored by hindpaw PRV infection in mice. Interestingly, this increased ATF3 in PRV-infected mice is preferentially detected in small-diameter sensory neurons of the DRG. These previous studies, combined with our data, indicate that the source of IL-6 may be the injured DRG neurons or surrounding satellite glia cells, which can then activate the HPA axis and corticosterone-dependent stress response (KAPCALA et al., 1995; Späth-Schwalbe et al., 1994) to cause the thymic atrophy (Ansari and Liu, 2017; Carbajosa et al., 2017; Sempowski et al., 2000). Notably, anti-IL-6 treatment did not have any substantial effect on the development of “mad itch” behaviors or appearance of skin lesions but rescues the thymic T cell development defects to a large extent. Thus, a scenario emerges where injured sensory neurons cause an increase in serum IL-6, activation of the hypothalamus and increased serum corticosterone levels which may lead to loss of thymic architecture and impaired development of T cells, which are important for anti-PRV immunity (Bianchi et al., 1998).

Our data uncovers critical strategies employed by PRV to transport anterogradely along axons by hijacking DDX3X. Reducing the levels of DDX3X leads to the death of sensory neurons. The loss of sensory neurons has three major effects which further facilitates the spread of the virus: 1) reduced pain allows the infected host to feel unimpaired, 2) an addictive “mad itch” causes tissue damage exposing the virus to potential new hosts, and 3) thymic atrophy and reduced peripheral T cell numbers to divert anti-viral immunity. Our results identify molecular and cellular mechanisms of how a virus can hijack an essential host protein for its intricate life cycle and provide new tools and insights on one of the most idiosyncratic medical mysteries - neuropathic “mad itch”.

## Acknowledgements.

This work was supported by the National key research and development plan, Program Intergovernmental cooperation, 2021YFE0193400 (SR, MJ and SJFC); by the OeAD China- Austria Cooperation Research Grant (SJFC, MJ and SR); by the Swedish Research Council (Swedish research Council 2018-05766) (JMP and SJFC); by the National Institute of Neurological disorders and Stroke (NINDS) grants R01 NS33506 and NS060699 (KL); by the Swiss National Science Foundation P400PM_194473, the Bruno Bloch Foundation (OHA); by the Austrian Science Fund (FWF), SFB F6103 and W1261 (TD); by the Austrian Federal Ministry of Education, Science and Research, the Austrian Academy of Sciences, and the City of Vienna, and grants from the T. von Zastrow Foundation, the Austrian Science Fund (FWF), Wittgenstein award (Z 271-B19), and a Canada 150 Research Chairs Program (F18-01336) (all to JMP). We acknowledge the Vienna BioCenter Core Facilities (VBCF) for their histopathology work.

## Author Contributions

SJFC, together with OK and JMP, conceived and designed the study. PRV-related infection experiments were performed by RS, LK and OK. All other experiments were coordinated by SJFC. All experiments were performed by SJFC with the following exceptions: MAT and DC helped with mouse behavior; JS performed mass spectrometry; MN and SR analyzed RNA sequencing data; JL performed MRI studies; SR, AH and OHA performed ELISA; TP and TD previously generated the *Ddx3x*-floxed animal; MJ, LB and AB provided DDX3X inhibitors. SJFC, together with OK and JMP, wrote the manuscript with input from all authors.

## Methods

### Cell lines and virus strains

The porcine kidney (PK15) epithelial cells and African green monkey kidney epithelial cells (Vero) were purchased from the American Type Culture Collection (ATCC) and were used to grow and titer PRV and HSV-1, respectively. Cells were maintained in Dulbecco’s modified Eagle medium supplemented with 10% fetal bovine serum, 1% penicillin and streptomycin. PRV Becker is a wild-type strain (Platt et al., 1980) virus: thermal sensitivity and rabbit virulence markers. PRV Bartha is an attenuated vaccine strain (Bartha, 1969). PRV-180 expresses mRFP-VP26 in a PRV-Becker background (del Rio et al., 2005). PRV 959, expressing mNeonGreen-VP26 in a PRV-Becker background, was constructed by Jens B Bosse, Ian B Hogue, and Lynn W Enquist (Koyuncu et al., 2015). Fluorescent HSV-1 recombinants were derived from HSV-1 strain 17. HSV-1 OK14 expresses an mRFP fusion to the capsid protein VP26 (Song et al., mBio2016).

### Neuronal cultures

Superior cervical ganglia (SCG) and dorsal root ganglia (DRG) were isolated from E16-17 Sprague-Dawley rat embryos (Hilltop Labs, Inc., Scottsdale, PA) and neurons were cultured in tri-chambers as described (Ch’ng and Enquist, 2005). All animal work was performed in accordance with the Princeton Institutional Animal Care and Use Committee (protocols 1947-13 and 1851-14). Multiwell dishes (Falcon), 35-mm plastic tissue culture dishes (Falcon) or optical plastic dishes (Ibidi) were coated with 500 µg/ml of poly-DL-ornithine (Sigma Aldrich) and 10 µg/ml of natural mouse laminin (Life Technologies). To prepare compartmented neuronal cultures, two sets of evenly spaced parallel grooves were etched on the dishes before a silicone grease-coated tri-chamber (Tyler Research) was placed. SCGs were trypsinized and triturated before plating in the Soma (S) compartment. Neurons were maintained in neurobasal medium (Gibco) supplemented with 100 ng/ml nerve growth factor 2.5S (Invitrogen), 2% B27 (Gibco) and 1% penicillin and streptomycin with 2 mM glutamine (Life Technologies). Two days after plating, 1 mM cytosine-D-arabinofuranoside (Sigma-Aldrich) was added to eliminate non-neuronal dividing cells. Neurons were cultured for 14-21 days.

### Viral infection and inhibitor treatment in compartmented neuronal cultures

For all experiments in tri-chambers, 1% methylcellulose in neuronal medium was added in the M (Middle) compartment before infection. Neuronal infections were done using 10^6^ or 10^4^ pfu (plaque forming units) of PRV. DiO was added at 2.5 μg/mL (Life Technologies) to the N (neurite/epithelial) compartment at 1 hpi. DDX3X inhibitor was added to the cell body-compartment media 1 hpi at 20 µM concentration. For titer assays, contents of the cell body or neurite/epithelial compartments were collected by scraping the dish with a pipette tip. All viral titers were determined on PK15 cells by serial dilution and are expressed in pfu/ml.

### Microscopy

Neuronal cultures were imaged using a Nikon Ti-Eclipse inverted epifluorescence microscope. Tiled images of the entire S compartment were captured using a Cool Snap ES2 camera (Photometrics) at 4x magnification. During overnight imaging, neuron cultures were kept in a humidified stage-top incubator at 37°C with 5% CO_2_. The percentage of infected cell bodies and moving fluorescent particles were calculated manually using NIS-Elements imaging software (Nikon). For fixed samples, images were captured using the Cool Snap ES2 camera at 60x. All images and movies were assembled for publication using NIS software (Nikon) or ImageJ. For comparative analysis, fluorescence intensity, exposure time, and other parameters were consistent for all conditions in the same experiment.

### Reagents

Anti-mouse IL-6 (Bio-X-Cell, BE0046) and anti-mouse Ly6G (Bio-X-Cell, BE0075-1); G-CSF ELISA kit (R&D Systems, MCS00); Corticosterone ELISA (R&D Systems, KGE009); IL-6 ELISA (Biolegend, 431304); Histamine ELISA (Beckman Coulter, IM2015).

### Western blot analysis

Dissociated neurons in multiwall dishes or tissues were lysed in radioimmunoprecipitation assay (RIPA) buffer without sodium deoxycholate, supplemented with 1 mM dithiothreitol (DTT) and protease inhibitor cocktail (Sigma-Aldrich). Lysates were incubated on ice for 30 min, briefly sonicated, and centrifuged at 11,000 rpm at 4°C. Cleared supernatants were transferred into new tubes and equal volumes were mixed with 5x Laemmli buffer. Samples were heated at 95°C for 5 min before being resolved by 8-10% SDS polyacrylamide gel electrophoresis. Gels were transferred to nitrocellulose membranes (Whatman) using semi-dry transfer (Biorad). After transfer, membranes were blocked in 5% non-fat dry milk in tris-buffered saline (TBS) solution for 1 h at room temperature. Immunoblots were performed using primary and secondary antibodies in 1% milk-TBS solution. Membranes were incubated with chemiluminescent substrates (Supersignal West Pico or Dura, Thermo scientific). Protein bands were visualized by exposure on HyBlot CL (Denville scientific) blue X-ray films. Immunoblots were performed using the following primary antibodies: anti-β-actin (1:5000) (Sigma), mouse monoclonal anti-VP5 (1:2000) (gift of H. Rziha), rabbit polyclonal anti-Us9 (1∶1000) (Brideau, Banfield, Enquist, 1998). Primary antibodies were detected using horseradish peroxidase-conjugated secondary mouse or rabbit antibodies (1:5000) (KPL). Other antibodies used for protein detection included: anti- DDX3X (Bethyl; A300-474A), anti-KIF1A (BD Bioscience; 612094), anti-beta-tubulin III (Abcam; ab18207), and anti-beta Actin (Sigma; A2228).

### Immunofluorescence staining of cultured neurons

Neurons were either cultured in chambered optical plastic dishes (Ibidi) or in multiwell dishes with glass coverslips. Fixations were done using 4% paraformaldehyde (Electron Microscopy Sciences) in phosphate-buffered saline (PBS) for 10 min and permeablized with ice-cold methanol for 10 min at −20°C. Samples were then incubated in PBS containing 3% bovine serum albumin (BSA) (Sigma-Aldrich) for 1 h prior to staining with primary and secondary antibodies and DAPI diluted in 3% BSA in PBS. Staining was performed using an anti-DDX3X antibody (1:500, santa cruz; sc- 365768) detected by an Alexa Fluor 488-conjugated secondary antibody against rabbit (1:500) (Life Technologies) and counterstained to detect nuclei using DAPI (1:1000). Teflon tri-chambers were removed before mounting with Aqua-Poly/Mount (Polysciences) and covering with glass slips (Fisher Scientific).

### Mice

*Ddx3x* floxed animals were previously generated and described by our laboratory (Szappanos et al., 2018). The following *Cre*-expressing lines were crossed to *Ddx3x* floxed animals to generate tissue specific conditional KO animals: *NaV1.8-Cre* line (Agarwal et al., 2004) and tamoxifen-inducible *Brn3a-Ert-Cre* mice (O’Donovan et al., 2014). The following reporter mice and ablation mice were used in the study: lox-STOP-lox transgenic *YFP*-reporter mice (Srinivas et al., 2001), conditional lox-STOP-lox transgenic diphtheria toxin (*DTx*) mice, conditional lox-STOP-lox transgenic diphtheria receptor (*DTR*) mice (Buch et al., 2005). The following *Cre*-expressing immune cell type lines were used to dissect immune involvement: *Mrp8-Cre* (Passegué et al., 2004), *Cpa3-Cre* (Lilla et al., 2011), *Rag1-Cre* (McCormack et al., 2003). All animal experiments were approved by the Austrian Animal Care and Use Committee.

### Behavioral assays

All assays were approved by the Austrian Animal Care and Use Committee and conducted in a blinded fashion in a quiet room (temperature 22±1°C) from 9 AM to 6 PM.

Mice were housed with their littermates (2 to 5 mice per cage based on the litters) with food and water ad libitum. All animals were maintained under the same conditions (22±1°C, 50% relative humidity, 12-h light/dark cycle). For behavioral experiments involving transgenic mice, randomization was achieved through the breeding: at the time of weaning mice were separated based on their sex and placed in their new home cage. Only cages with a mixed representation of genetically modified mice and their respective littermates were used for behavioral experiments. All experiments used at least 2 independent litters and were performed at least twice.

### Determination of pain responses/thresholds

Mice were transferred to a specialized mouse behavior room at least 1 week prior to experiments and housed at a 14 hours light - 10 hours dark cycle access to food and water *ad libitum*.

#### Contact heat pain (Hot plate test)

Mice were placed on a metallic plate heated to a set temperature within an acrylic container (Bioseb, France), and the latency for flinching, licking one of the hind paws, or jumping was measured. Mice were sequentially tested for 50°C and 52°C. One temperature was tested per day.

#### Radiant heat pain (Hargreaves test)

Before testing, each mouse was placed on an elevated glass surface and habituated in their individual cage for 1 hour. Then a radiant heat source (Ugo Basile plantar test model 7371/7372) was targeted at the left hind paw and the latency to withdraw was measured.

#### Capsaicin response

20 microliters of 2 μg capsaicin (Sigma, M2028) diluted in 1% DMSO and saline were injected into the plantar surface of the hind paw and the mouse was placed onto a room temperature surface within a container. The time spent licking, biting or lifting the paw was measured.

#### Pin prick

Pinprick was assessed using an Austerlitz insect pin (size, 000; Fine Science Tools). This pin was gently applied to the lateral plantar surface of the paw without moving the paw and at an intensity sufficient to indent but not penetrate the skin. Positive responses from 10 applications were identified by rapid paw withdrawal immediately after stimulation.

### PRV infection of mice

The protocol used for the footpad inoculation experiments was previously described (Laval et al., 2019). Briefly, mice were anesthetized with 1–3% isoflurane gas and the right hind footpad, between the heel and walking pads, was gently abraded about 20 times with an emery board until the stratum corneum was removed. A 20-μl droplet of virus inoculum containing 8X10^6^ plaque-forming unit (PFU) of PRV-Becker, resuspended in medium (Dulbecco modified Eagle medium, 2% fetal calf serum and antibiotics) (Hyclone, GE Healthcare life sciences), was applied onto the abraded area of the skin. Mock-inoculations (medium only) were carried out in parallel. The inoculum was gently rubbed 5 to 10 times with the shaft of an 18-gauge hypodermic needle to facilitate adsorption of the virus. The mice were kept under anesthesia for 30 minutes (min) until the abraded footpad was dry and then the animals were placed in separate cages for further analysis

### Immunostaining of DRG tissue

L4-L5 DRG tissue were extracted, fixed in 4% paraformaldehyde dissolved in PBS, cryoprotected in 30% sucrose, and frozen in OCT (Tissue-Tek). Ten-μm thick cryosections were blocked with 1% bovine serum albumin (Sigma-Aldrich)/ 0.1%Triton X-100 in 0.1 M phosphate buffered saline (PBS) and then incubated with primary antibodies overnight at 4°C. After 3 washes in PBS for 10 minutes each, sections were incubated with secondary antibodies for 1 hour at room temperature, washed 3 times in PBS (10 minutes each) and mounted using Dako mounting medium (S3023). The following primary antibodies were used: Isolectin B4 (IB4) DyLight 594-conjugated (Vector laboratories FL- 1207); CGRP (Millipore; PC205L), DDX3X (santa cruz; sc-365768); NF200 (Abcam; ab4680); Parvalbumin (Swant; PV-25). Secondary antibodies used: Alexa Fluor 555 anti-rabbit (Invitrogen; A32794), Alexa Fluor 555 anti-mouse (Invitrogen; A32773); Alex Fluor 488 anti-rabbit (Invitrogen; A21206); beta-tublin-488 (BD Pharmingen; 560338).

### Serum cytokine analysis

To measure proinflammatory cytokine levels in the serum of mice, ProcartaPlex Immunoassays (Thermo Fisher Scientific) were used. Blood was collected in a BD Microtainer blood collection tube to avoid coagulation. Subsequently, blood was centrifuged at 6000xg for 3 minutes and serum was collected. Samples were measured in duplicates following the manufacturer’s instructions. The fluorescence intensity for each cytokine of each sample was read in a Luminex analyzer.

### Manganese-enhanced magnetic resonance imaging (MEMRI)

Mice were imaged using a 15.2 T small animal MRI (Bruker BioSpec, Ettlingen, Germany) and 23 mm birdcage coil. Prior to imaging, animals were anesthetized with 4% isoflurane, and care was taken to adjust the isoflurane levels immediately so that respiration did not fall below 140 breaths per minute (bpm) at any time. During imaging, respiration was maintained between 140 and 160 bpm corresponding to about 1.4-1.8% isofluorane. To provide quantitative assessment of neuronal activation and Mn^2+^ accumulation in neurons we measured the longitudinal relaxation time (T_1_) before and after MnCl_2_ injections, since Mn^2+^ is known to reduce T_1_. Rapid acquisition with relaxation enhancement (RARE) sequence with variable repetition time VTR=(300, 500, 750, 1500, 3000, 6000) ms, TE=6 ms, RARE factor=2, matrix 256X256, FOV 20X20 cm, NEX=1, and total imaging time 25 min was used to quantify T_1_. Prior to MnCl_2_ injection, a baseline scan was performed. Mice were *i.p*. injected with 50 mg/kg MnCl_2_ for two times (two hours apart) and imaged 24 h after the first injection to determine changes in T_1_ values. Images were realigned and T_1_ valuescalculated using ImageJ. Relaxivity rates (R_1_) were calculated as R_1_=1/T_1_. Calculated R_1_ maps were threshold based on R1 values compared with baseline scans. We took 40% increase in R_1_ values to be strong significant indicator of Mn accumulation as a measure of neuronal activation. Regions that had > 40% R_1_ increase were overlaid over anatomical brain images (magnitude images from T_1_ quantification).

### Flow cytometry

Thymi and spleens were extracted, and single cell suspensions were made. The dissociated cells were passed through a 70-μm cell strainer and then washed with 10 ml cold FACS buffer. The cells were stained with directly fluorescence labelled antibodies specific to CD45, CD4, CD8, B220, TCRbeta, CD11b, Gr-1, Ly-6G. Unspecific staining was blocked using an anti-mouse CD16/CD32 Fc block (553142, BD Biosciences, 1:100) and DAPI (D1306, Thermo Fisher Scientific, 1:500 from a 5 mg/ml stock). All reactions were performed in FACS buffer and incubated for 20 min at 4°C. Cells were analyzed on a BD LSR Fortessa. All data were analyzed with FlowJo v10.0.8r1.

### Immunoprecipitations

An equal amount of protein lysate (sciatic nerve tissue) from wild type mice was incubated with anti-DDX3X or an isotype control IgG antibody for 2 hours at 4°C, followed by rotating incubation with 20 µl of protein G-Agarose beads (Sigma) for 1 hour. After several washes to remove unbound proteins, the immune complexes were analyzed by mass spectrometry and Western blot analyses.

### Mass spectroscopy

IPs were on-bead digested using trypsin and the recovered peptides were analyzed by tandem mass-spectrometry (MS/MS), using nano-reversed-phase-HPLC coupled to a QExactive HF-X instrument. MS/MS data were then searched against the Uniprot mouse proteome sequence database using MS Amanda (Dorfer et al., 2014).

### Histology

Dorsal skin from mice was fixed in 10% neutral buffered formalin (Sigma, HT501128), processed with a microwave hybrid tissue processor (LOGOS, Milestone Medical), embedded in paraffin, sectioned (Microm, HM 355), and stained with hematoxylin and eosin in an automated stainer (Microm HMS 740). Slides were evaluated and scored by a board-certified veterinary comparative pathologist. For histological analysis of the cortico-medullary organization of the thymus, 2.5-μm paraffin-sections were stained with haematoxylin and eosin (H&E). Slides were then scanned on a Mirax Scanner (Zeiss).

### RNA sequencing and analysis

DRG tissue was extracted 2 days after tamoxifen treatment ended in *Ddx3x^BRN3A^* and littermate control mice. RNA was extracted and processed for RNA-seq experiments. Data were analysed using genome and UCSC gene annotation provided in the iGenomes UCSC mm 10bundle (https://support.illumina.com/sequencing/sequencing_software/igenome.html). RNA-seq reads were first trimmed using trimgalore v0.5.0 and reads mapping to abundant sequences included in the iGenomes UCSC mm10 bundle (mouse rDNA, mouse mitochondrial chromosome, phiX174 genome, adapter) were removed using bowtie2 v2.3.4.1 alignment. Remaining reads were aligned to the mm10 genome using star v2.6.0c and reads in genes were counted with feature Counts (subread v1.6.2) and parameter -s2. Differential gene expression analyses on raw counts were performed using DESeq2 v1.18.1. Gene set over- representation analysis of differentially expressed genes was conducted using clusterprofiler v3.6.0 in R v3.4.1.

### Chemicals and reagents

The following reagents were used in this study: diphtheria toxin (Sigma, D0564), tamoxifen (Sigma, T5648), naloxone (Sigma, N7758), anti-mouse Ly-6G antibody (BioxCell, BE0075-1), anti-mouse IL-6 antibody (BioXCell, BE0046), anti-mouse IL-6 ELISA kit (Sigma, RAB0309), anti-mouse G-CSF ELISA kit (MCS00, R&D Systems), IgE ELISA kit (company?), and a histamine ELISA kit (company?). The DDX3X inhibitor 16d was synthesized by co-author Lorenzo Botta (Brai et al., 2016).

### Statistical analysis

All values are expressed as means +/- s.e.m. Details of the statistical tests used are stated in each figure legend. Briefly, Student’s t-test was used to compare between two groups. One-way ANOVA followed by Dunnett’s post-hoc test for multiple comparisons was used for analysis between multiple groups. 2-way ANOVA was used to compare two groups over time. In all tests P ≤ 0.05 was considered significant.

## References

Agarwal, N., Offermanns, S., and Kuner, R. (2004). Conditional Gene Deletion in Primary Nociceptive Neurons of Trigeminal Ganglia and Dorsal Root Ganglia. Genesis 38.

Albano, F., Vecchio, E., Renna, M., Iaccino, E., Mimmi, S., Caiazza, C., Arcucci, A., Avagliano, A., Pagliara, V., Donato, G., et al. (2019). Insights into thymus development and viral thymic infections. Viruses 11.

Alcami, A., and Koszinowski, U.H. (2000). Viral mechanisms of immune evasion. Trends Microbiol. 8.

Ansari, A.R., and Liu, H. (2017). Acute Thymic Involution and Mechanisms for Recovery. Arch. Immunol. Ther. Exp. (Warsz). 65.

Babic, N., Mettenleiter, T.C., Ugolini, G., Flamand, A., and Coulon, P. (1994). Propagation of Pseudorabies Virus in the Nervous System of the Mouse after Intranasal Inoculation. Virology 204.

Barr, A.R., and Gergely, F. (2008). MCAK-Independent Functions of ch-Tog/XMAP215 in Microtubule Plus-End Dynamics. Mol. Cell. Biol. 28.

Bergasa, N.V., Ailing, D.W., Talbot, T.L., Swain, M.G., Yurdaydin, C., Turner, M.L., Schmitt, J.M., Walker, E.C., and Jones, E.A. (1995). Effects of naloxone infusions in patients with the pruritus of cholestasis: A double-blind, randomized, controlled trial. Ann. Intern. Med. 123.

Berridge, K.C., and Kringelbach, M.L. (2015). Pleasure Systems in the Brain. Neuron 86.

Bianchi, A.T.J., Moonen-Leusen, H.W.M., Van Milligen, F.J., Savelkoul, H.F.J., Zwart, R.J., and Kimman, T.G. (1998). A mouse model to study immunity against pseudorabies virus infection: Significance of CD4+ and CD8+ cells in protective immunity. Vaccine 16.

Borriello, F., Iannone, R., and Marone, G. (2017). Histamine release from mast cells and basophils. In Handbook of Experimental Pharmacology, p.

Brai, A., Fazi, R., Tintori, C., Zamperini, C., Bugli, F., Sanguinetti, M., Stigliano, E., Esté, J., Badia, R., Franco, S., et al. (2016). Human DDX3 protein is a valuable target to develop broad spectrum antiviral agents. Proc. Natl. Acad. Sci. U. S. A. 113.

Brai, A., Martelli, F., Riva, V., Garbelli, A., Fazi, R., Zamperini, C., Pollutri, A., Falsitta, L., Ronzini, S., Maccari, L., et al. (2019). DDX3X Helicase Inhibitors as a New Strategy to Fight the West Nile Virus Infection. J. Med. Chem. 62.

Brideau, A.D., Card, J.P., and Enquist, L.W. (2000). Role of Pseudorabies Virus Us9, a Type II Membrane Protein, in Infection of Tissue Culture Cells and the Rat Nervous System. J. Virol. 74.

Brittle, E.E., Reynolds, A.E., and Enquist, L.W. (2004). Two Modes of Pseudorabies Virus Neuroinvasion and Lethality in Mice. J. Virol. 78.

Brown, C.R., Blaho, V.A., and Loiacono, C.M. (2004). Treatment of mice with the neutrophil-depleting antibody RB6-8C5 results in early development of experimental lyme arthritis via the recruitment of Gr-1- polymorphonuclear leukocyte-like cells. Infect. Immun. 72.

Buch, T., Heppner, F.L., Tertilt, C., Heinen, T.J.A.J., Kremer, M., Wunderlich, F.T., Jung, S., and Waisman, A. (2005). A Cre-inducible diphtheria toxin receptor mediates cell lineage ablation after toxin administration. Nat. Methods 2.

Buddenkotte, J., Maurer, M., and Steinhoff, M. (2010). Histamine and antihistamines in atopic dermatitis. Adv. Exp. Med. Biol. 709.

Carbajosa, S., Gea, S., Chillón-Marinas, C., Poveda, C., Maza, M. del C., Fresno, M., and Gironès, N. (2017). Altered bone marrow lymphopoiesis and interleukin-6-dependent inhibition of thymocyte differentiation contribute to thymic atrophy during Trypanosoma cruzi infection. Oncotarget 8.

Ch’ng, T.H., and Enquist, L.W. (2005). Neuron-to-Cell Spread of Pseudorabies Virus in a Compartmented Neuronal Culture System. J. Virol. 79.

Chen, X.J., and Sun, Y.G. (2020). Central circuit mechanisms of itch. Nat. Commun. 11.

Chen, C.Y., Chan, C.H., Chen, C.M., Tsai, Y.S., Tsai, T.Y., Lee, Y.H.W., and You, L.R. (2016a). Targeted inactivation of murine Ddx3x: Essential roles of Ddx3x in placentation and embryogenesis. Hum. Mol. Genet. 25.

Chen, H.H., Yu, H.I., and Tarn, W.Y. (2016b). DDX3 modulates neurite development via translationally activating an RNA regulon involved in Rac1 activation. J. Neurosci. 36.

Chiba, K., Takahashi, H., Chen, M., Obinata, H., Arai, S., Hashimoto, K., Oda, T., McKenney, R.J., and Niwa, S. (2019). Disease-associated mutations hyperactivate KIF1A motility and anterograde axonal transport of synaptic vesicle precursors. Proc. Natl. Acad. Sci. U. S. A. 116.

Ciccosanti, F., Di Rienzo, M., Romagnoli, A., Colavita, F., Refolo, G., Castilletti, C., Agrati, C., Brai, A., Manetti, F., Botta, L., et al. (2021). Proteomic analysis identifies the RNA helicase DDX3X as a host target against SARS-CoV-2 infection. Antiviral Res. 190.

Damann, N., Klopfleisch, R., Rothermel, M., Doerner, J.F., Mettenleiter, T.C., Hatt, H., and Wetzel, C. (2006). Neuronal pathways of viral invasion in mice after intranasal inoculation of pseudorabies virus PrV-9112C2 expressing bovine herpesvirus 1 glycoprotein B. J. Neurovirol. 12.

Ditton, H.J., Zimmer, J., Kamp, C., Rajpert-De Meyts, E., and Vogt, P.H. (2004). The AZFa gene DBY (DDX3Y) is widely transcribed but the protein is limited to the male germ cells by translation control. Hum. Mol. Genet. 13.

Dong, X., and Dong, X. (2018). Peripheral and Central Mechanisms of Itch. Neuron 98.

Dorfer, V., Pichler, P., Stranzl, T., Stadlmann, J., Taus, T., Winkler, S., and Mechtler, K. (2014). MS Amanda, a universal identification algorithm optimized for high accuracy tandem mass spectra. J. Proteome Res. 13.

Dubový, P., Brázda, V., Klusáková, I., and Hradilová-Svíženská, I. (2013). Bilateral elevation of interleukin-6 protein and mRNA in both lumbar and cervical dorsal root ganglia following unilateral chronic compression injury of the sciatic nerve. J. Neuroinflammation 10.

DuRaine, G., and Johnson, D.C. (2021). Anterograde transport of α-herpesviruses in neuronal axons. Virology 559.

Esancy, K., Condon, L., Feng, J., Kimball, C., Curtright, A., and Dhaka, A. (2018). A zebrafish and mouse model for selective pruritus via direct activation of TRPA1. Elife 7.

Fairman-Williams, M.E., Guenther, U.P., and Jankowsky, E. (2010). SF1 and SF2 helicases: Family matters. Curr. Opin. Struct. Biol. 20.

Field, H.J., and Hill, T.J. (1974). The pathogenesis of pseudorabies in mice following peripheral inoculation. J. Gen. Virol. 23.

Fitzek, S., Baumgärtner, U., Marx, J., Joachimski, F., Axer, H., Witte, O.W., and Fitzek, C. (2006). Chapter 15 Pain and itch in Wallenberg’s syndrome: anatomical-functional correlations. Suppl. Clin. Neurophysiol. 58.

Fukuyama, R., and Rapoport, S.I. (1995). Brain-specific expression of human microtubule-associated protein 1A (MAP1A) gene and its assignment to human chromosome 15. J. Neurosci. Res. 40.

Fuller-Pace, F. V. (2006). DExD/H box RNA helicases: Multifunctional proteins with important roles in transcriptional regulation. Nucleic Acids Res. 34.

Gautron, L., Sakata, I., Udit, S., Zigman, J.M., Wood, J.N., and Elmquist, J.K. (2011). Genetic tracing of Nav1.8-expressing vagal afferents in the mouse. J. Comp. Neurol. 519.

Gimeno, I.M., Witter, R.L., Cortes, A.L., and Reed, W.M. (2011). Replication ability of three highly protective Marek’s disease vaccines: Implications in lymphoid organ atrophy and protection. Avian Pathol. 40.

Gulland, F.M.D., Lowenstine, L.J., Lapointe, J.M., Spraker, T., and King, D.P. (1997). Herpesvirus infection in stranded pacific harbor seals of coastal California. J. Wildl. Dis. 33.

Guo, Z., Chen, X.X., and Zhang, G. (2021). Human PRV Infection in China: An Alarm to Accelerate Eradication of PRV in Domestic Pigs. Virol. Sin. 36.

Halpain, S., and Dehmelt, L. (2006). The MAP1 family of microtubule-associated proteins. Genome Biol. 7.

Hao, Z., and Rajewsky, K. (2001). Homeostasis of peripheral B cells in the absence of B cell influx from the bone marrow. J. Exp. Med. 194.

Heath, R.G. (1963). ELECTRICAL SELF-STIMULATION OF THE BRAIN IN MAN. Am. J. Psychiatry 120.

Hu, D., Al-Shalan, H.A.M., Shi, Z., Wang, P., Wu, Y., Nicholls, P.K., Greene, W.K., and Ma, B. (2020). Distribution of nerve fibers and nerve-immune cell association in mouse spleen revealed by immunofluorescent staining. Sci. Rep. 10.

Huang, H., Koyuncu, O.O., and Enquist, L.W. (2020). Pseudorabies Virus Infection Accelerates Degradation of the Kinesin-3 Motor KIF1A. J. Virol. 94.

Huang, S., Ziegler, C.G.K., Austin, J., Mannoun, N., Vukovic, M., Ordovas-Montanes, J., Shalek, A.K., and von Andrian, U.H. (2021). Lymph nodes are innervated by a unique population of sensory neurons with immunomodulatory potential. Cell 184.

Husak, P.J., Kuo, T., and Enquist, L.W. (2000). Pseudorabies Virus Membrane Proteins gI and gE Facilitate Anterograde Spread of Infection in Projection- Specific Neurons in the Rat. J. Virol. 74.

Ivanova, A., Signore, M., Caro, N., Greene, N.D.E., Copp, A.J., and Martinez-Barbera, J.P. (2005). In vivo genetic ablation by Cre-mediated expression of diphtheria toxin fragment A. Genesis 43.

Ju, T., and Yosipovitch, G. (2020). Neuropathic pruritus associated with brain disorders. Itch 5.

Kapcala, L.P., Chautard, T., and Eskay, R.L. (1995). The Protective Role of the Hypothalamic-Pituitary-Adrenal Axis against Lethality Produced by Immune, Infectious, and Inflammatory Stress. Ann. N. Y. Acad. Sci. 771.

Kesavardhana, S., Samir, P., Zheng, M., Subbarao Malireddi, R.K., Karki, R., Sharma, B.R., Place, D.E., Briard, B., Vogel, P., and Kanneganti, T.D. (2021). DDX3X coordinates host defense against influenza virus by activating the NLRP3 inflammasome and type I interferon response. J. Biol. Chem. 296.

Khadivjam, B., Stegen, C., Hogue-Racine, M.-A., El Bilali, N., Döhner, K., Sodeik, B., and Lippé, R. (2017). The ATP-Dependent RNA Helicase DDX3X Modulates Herpes Simplex Virus 1 Gene Expression. J. Virol. 91.

Ko, M.C. (2015). Neuraxial opioid-induced itch and its pharmacological antagonism. Handb. Exp. Pharmacol. 226.

Koyuncu, O.O., Song, R., Greco, T.M., Cristea, I.M., and Enquist, L.W. (2015). The number of alphaherpesvirus particles infecting axons and the axonal protein repertoire determines the outcome of neuronal infection. MBio 6.

Koyuncu, O.O., Enquist, L.W., and Engel, E.A. (2020). Invasion of the nervous system. Curr. Issues Mol. Biol. 41.

Kramer, T., Greco, T.M., Enquist, L.W., and Cristea, I.M. (2011). Proteomic Characterization of Pseudorabies Virus Extracellular Virions. J. Virol. 85.

Kramer, T., Greco, T.M., Taylor, M.P., Ambrosini, A.E., Cristea, I.M., and Enquist, L.W. (2012). Kinesin-3 mediates axonal sorting and directional transport of alphaherpesvirus particles in neurons. Cell Host Microbe 12.

Kratchmarov, R., Taylor, M.P., and Enquist, L.W. (2012). Making the case: Married versus Separate models of alphaherpes virus anterograde transport in axons. Rev. Med. Virol. 22.

Langan, S.M., Irvine, A.D., and Weidinger, S. (2020). Atopic dermatitis. Lancet 396.

Laval, K., and Enquist, L.W. (2020). The neuropathic itch caused by pseudorabies virus. Pathogens 9.

Laval, K., Vernejoul, J.B., Van Cleemput, J., Koyuncu, O.O., and Enquist, L.W. (2018). Virulent Pseudorabies Virus Infection Induces a Specific and Lethal Systemic Inflammatory Response in Mice. J. Virol. 92.

Laval, K., Van Cleemput, J., Vernejoul, J.B., and Enquist, L.W. (2019). Alphaherpesvirus infection of mice primes PNS neurons to an inflammatory state regulated by TLR2 and type i IFN signaling. PLoS Pathog. 15.

Li, J., Zhang, W., Guo, Z., Wu, S., Jan, L.Y., and Jan, Y.N. (2016). A defensive kicking behavior in response to mechanical stimuli mediated by Drosophila wing margin bristles. J. Neurosci. 36.

Lilla, J.N., Chen, C.G., Mukai, K., BenBarak, M.J., Franco, C.B., Kalesnikoff, J., Yu, M., Tsai, M., Piliponsky, A.M., and Galli, S.J. (2011). Reduced mast cell and basophil numbers and function in Cpa3-Cre; Mcl-1 fl/fl mice. Blood 118.

Lin, Y.J., and Koretsky, A.P. (1997). Manganese ion enhances T1-weighted MRI during brain activation: An approach to direct imaging of brain function. Magn. Reson. Med. 38.

Linder, P., and Jankowsky, E. (2011). From unwinding to clamping ĝ€" the DEAD box RNA helicase family. Nat. Rev. Mol. Cell Biol. 12.

Lipman, Z.M., and Yosipovitch, G. (2021). Substance use disorders and chronic itch. J. Am. Acad. Dermatol. 84.

Lipshetz, B., Khasabov, S.G., Truong, H., Netoff, T.I., Simone, D.A., and Giesler, G.J. (2018). Responses of thalamic neurons to itch-and pain-producing stimuli in rats. J. Neurophysiol. 120.

Loret, S., Guay, G., and Lippé, R. (2008). Comprehensive Characterization of Extracellular Herpes Simplex Virus Type 1 Virions. J. Virol. 82.

Mack, M.R., and Kim, B.S. (2018). The Itch–Scratch Cycle: A Neuroimmune Perspective. Trends Immunol. 39.

McCormack, M.P., Forster, A., Drynan, L., Pannell, R., and Rabbitts, T.H. (2003). The LMO2 T-Cell Oncogene Is Activated via Chromosomal Translocations or Retroviral Insertion during Gene Therapy but Has No Mandatory Role in Normal T-Cell Development . Mol. Cell. Biol. 23.

McVeigh, K.A., Adams, M., Harrad, R., and Ford, R. (2018). Periocular manifestations of trigeminal trophic syndrome: A case series and literature review. Orbit (London) 37.

Mettenleiter, T.C. (2000). Aujeszky’s disease (pseudorabies) virus: The virus and molecular pathogenesis - State of the art, June 1999. Vet. Res. 31.

Misery, L., Bodere, C., Genestet, S., Zagnoli, F., and Marcorelles, P. (2014). Small-fibre neuropathies and skin: News and perspectives for dermatologists. Eur. J. Dermatology 24.

Narita, M., and Ishi, M. (2006). Brain lesions in pigs dually infected with porcine reproductive and respiratory syndrome virus and pseudorabies virus. J. Comp. Pathol. 134.

Nelson, C., Mrozowich, T., Gemmill, D.L., Park, S.M., and Patel, T.R. (2021). Human ddx3x unwinds Japanese encephalitis and zika viral 5՛ terminal regions. Int. J. Mol. Sci. 22.

O’Donovan, K.J., Ma, K., Guo, H., Wang, C., Sun, F., Han, S.B., Kim, H., Wong, J.K., Charron, J., Zou, H., et al. (2014). B-RAF kinase drives developmental axon growth and promotes axon regeneration in the injured mature CNS. J. Exp. Med.

Oaklander, A.L. (2011). Neuropathic itch. Semin. Cutan. Med. Surg. 30.

Oaklander, A.L., Cohen, S.P., and Raju, S.V.Y. (2002). Intractable postherpetic itch and cutaneous deafferentation after facial shingles. Pain 96.

Oaklander, A.L., Bowsher, D., Galer, B., Haanpää, M., and Jensen, M.P. (2003). Herpes zoster itch: Preliminary epidemiologic data. J. Pain 4.

Ogawa, N., Kawai, H., Terashima, T., Kojima, H., Oka, K., Chan, L., and Maegawa, H. (2014). Gene therapy for neuropathic pain by silencing of TNF-α expression with lentiviral vectors targeting the dorsal root ganglion in mice. PLoS One 9.

Okada, Y., Yamazaki, H., Sekine-Aizawa, Y., and Hirokawa, N. (1995). The neuron-specific kinesin superfamily protein KIF1A is a uniqye monomeric motor for anterograde axonal transport of synaptic vesicle precursors. Cell 81.

Olds, J., and Milner, P. (1954). POSITIVE REINFORCEMENT PRODUCED BY ELECTRICAL STIMULATION OF SEPTAL AREA AND OTHER REGIONS OF RAT BRAIN. J. Comp. Physiol. Psychol. 47.

Otters, E.F.M., Tebbe-Gholami, M., and Weppner-Parren, L.J.M.T. (2014). Trigeminal trophic syndrome. Ned. Tijdschr. Voor Dermatologie En Venereol. 24.

Pantazis, C.B., and Aston-Jones, G. (2020). Lateral septum inhibition reduces motivation for cocaine: Reversal by diazepam. Addict. Biol. 25.

Passegué, E., Wagner, E.F., and Weissman, I.L. (2004). JunB deficiency leads to a myeloproliferative disorder arising from hematopoietic stem cells. Cell 119.

Patnaik, S., Imms, J.B., and Varadi, G. (2021). The neuropathic itch: Don’t scratch your head too hard! Clin. Case Reports 9.

Pereira, M.P., Mühl, S., Pogatzki-Zahn, E.M., Agelopoulos, K., and Ständer, S. (2016). Intraepidermal Nerve Fiber Density: Diagnostic and Therapeutic Relevance in the Management of Chronic Pruritus: a Review. Dermatol. Ther. (Heidelb). 6.

Platt, K.B., Maré, C.J., and Hinz, P.N. (1980). Differentiation of vaccine strains and field isolates of pseudorabies (Aujeszky’s disease) virus: Trypsin sensitivity and mouse virulence markers. Arch. Virol. 63.

Reich, A., and Szepietowski, J.C. (2010). Opioid-induced pruritus: An update. Clin. Exp. Dermatol. 35.

del Rio, T., Ch’ng, T.H., Flood, E.A., Gross, S.P., and Enquist, L.W. (2005). Heterogeneity of a Fluorescent Tegument Component in Single Pseudorabies Virus Virions and Enveloped Axonal Assemblies. J. Virol. 79.

Romero-Palomo, F., Risalde, M.A., Molina, V., Lauzi, S., Bautista, M.J., and Gómez-Villamandos, J.C. (2015). Characterization of thymus atrophy in calves with subclinical BVD challenged with BHV-1. Vet. Microbiol. 177.

Savino, W. (2006). The thymus is a common target organ in infectious diseases. PLoS Pathog. 2.

Sekiguchi, T., Iida, H., Fukumura, J., and Nishimoto, T. (2004). Human DDX3Y, the Y-encoded isoform of RNA helicase DDX3, rescues a hamster temperature-sensitive ET24 mutant cell line with a DDX3X mutation. Exp. Cell Res. 300.

Sempowski, G.D., Hale, L.P., Sundy, J.S., Massey, J.M., Koup, R.A., Douek, D.C., Patel, D.D., and Haynes, B.F. (2000). Leukemia Inhibitory Factor, Oncostatin M, IL-6, and Stem Cell Factor mRNA Expression in Human Thymus Increases with Age and Is Associated with Thymic Atrophy. J. Immunol. 164.

Soulat, D., Bürckstümmer, T., Westermayer, S., Goncalves, A., Bauch, A., Stefanovic, A., Hantschel, O., Bennett, K.L., Decker, T., and Superti-Furga, G. (2008). The DEAD-box helicase DDX3X is a critical component of the TANK-binding kinase 1-dependent innate immune response. EMBO J. 27.

Späth-Schwalbe, E., Born, J., Schrezenmeier, H., Bornstein, S.R., Stromeyer, P., Drechsler, S., Fehm, H.L., and Porzsolt, F. (1994). Interleukin-6 stimulates the hypothalamus-pituitary-adrenocortical axis in man. J. Clin. Endocrinol. Metab. 79.

Srinivas, S., Watanabe, T., Lin, C.S., William, C.M., Tanabe, Y., Jessell, T.M., and Costantini, F. (2001). Cre reporter strains produced by targeted insertion of EYFP and ECFP into the ROSA26 locus. BMC Dev. Biol. 1.

Steinhoff, M., Schmelz, M., Szabó, I.L., and Oaklander, A.L. (2018). Clinical presentation, management, and pathophysiology of neuropathic itch. Lancet Neurol. 17.

Strand, N., Mahdi, L., Schatman, M.E., Maloney, J., and Wie, C. (2021). Case study: Neuropathic itching following S3 and S4 dorsal root ganglion stimulator trial. J. Pain Res. 14.

Stumpf, A., and Ständer, S. (2013). Neuropathic itch: Diagnosis and management. Dermatol. Ther. 26.

Szappanos, D., Tschismarov, R., Perlot, T., Westermayer, S., Fischer, K., Platanitis, E., Kallinger, F., Novatchkova, M., Lassnig, C., Müller, M., et al. (2018). The RNA helicase DDX3X is an essential mediator of innate antimicrobial immunity. PLoS Pathog.

Tsherniak, A., Vazquez, F., Montgomery, P.G., Weir, B.A., Kryukov, G., Cowley, G.S., Gill, S., Harrington, W.F., Pantel, S., Krill-Burger, J.M., et al. (2017). Defining a Cancer Dependency Map. Cell 170.

Tsujino, H., Kondo, E., Fukuoka, T., Dai, Y., Tokunaga, A., Miki, K., Yonenobu, K., Ochi, T., and Noguchi, K. (2000). Activating transcription factor 3 (ATF3) induction by axotomy in sensory and motoneurons: A novel neuronal marker of nerve injury. Mol. Cell. Neurosci. 15.

van der Vaart, B., Franker, M.A.M., Kuijpers, M., Hua, S., Bouchet, B.P., Jiang, K., Grigoriev, I., Hoogenraad, C.C., and Akhmanova, A. (2012). Microtubule plus-end tracking proteins SLAIN1/2 and ch-TOG promote axonal development. J. Neurosci. 32.

Varnum, S.M., Streblow, D.N., Monroe, M.E., Smith, P., Auberry, K.J., Paša-Tolić, L., Wang, D., Camp, D.G., Rodland, K., Wiley, S., et al. (2004). Identification of Proteins in Human Cytomegalovirus (HCMV) Particles: the HCMV Proteome. J. Virol. 78.

Wang, D., Tao, X., Fei, M., Chen, J., Guo, W., Li, P., and Wang, J. (2020). Human encephalitis caused by pseudorabies virus infection: a case report. J. Neurovirol. 26.

Watanabe, K., Tanaka, J., Hatano, M., Nakanishi, I., Mukai, M., Nishikawa, J., Hayashi, S., and Kimura, N. (1984). Generalized neonatal herpes virus infection (cytomegalovirus or herpes virus type 1) comparative examination of loci attacked by two viruses. Pathol. Int. 34.

Wimalasena NK, et al. (2023). Nav1.7 gain-of-function mutation I228M triggers age-dependent nociceptive insensitivity and C-LTMR dysregulation. Exp Neurol.

Yamamoto, A., and Sugimoto, Y. (2010). Involvement of peripheral mu opioid receptors in scratching behavior in mice. Eur. J. Pharmacol. 649.

Yan, B., Wei, J.-J., Yuan, Y., Sun, R., Li, D., Luo, J., Liao, S.-J., Zhou, Y.-H., Shu, Y., Wang, Q., et al. (2013). IL-6 Cooperates with G-CSF To Induce Protumor Function of Neutrophils in Bone Marrow by Enhancing STAT3 Activation. J. Immunol. 190.

Yang, M., Card, J.P., Tirabassi, R.S., Miselis, R.R., and Enquist, L.W. (1999). Retrograde, Transneuronal Spread of Pseudorabies Virus in Defined Neuronal Circuitry of the Rat Brain Is Facilitated by gE Mutations That Reduce Virulence. J. Virol. 73.

Yu, C., Mannan, A.M., Yvone, G.M., Ross, K.N., Zhang, Y.L., Marton, M.A., Taylor, B.R., Crenshaw, A., Gould, J.Z., Tamayo, P., et al. (2016). High-throughput identification of genotype-specific cancer vulnerabilities in mixtures of barcoded tumor cell lines. Nat. Biotechnol. 34.

Zamboni, V., Armentano, M., Berto, G., Ciraolo, E., Ghigo, A., Garzotto, D., Umbach, A., Dicunto, F., Parmigiani, E., Boido, M., et al. (2018). Hyperactivity of Rac1-GTPase pathway impairs neuritogenesis of cortical neurons by altering actin dynamics. Sci. Rep. 8.

Zhang, P., Iwama, A., Datta, M.W., Darlington, G.J., Link, D.C., and Tenen, D.G. (1998). Upregulation of interleukin 6 and granulocyte colony-stimulating factor receptors by transcription factor CCAAT enhancer binding protein α (C/EBPα) is critical for granulopoiesis. J. Exp. Med. 188.

