## Supplemental data for "Pseudorabies virus hijacks DDX3X, initiating an addictive “mad itch” and immune suppression, to facilitate viral spread"

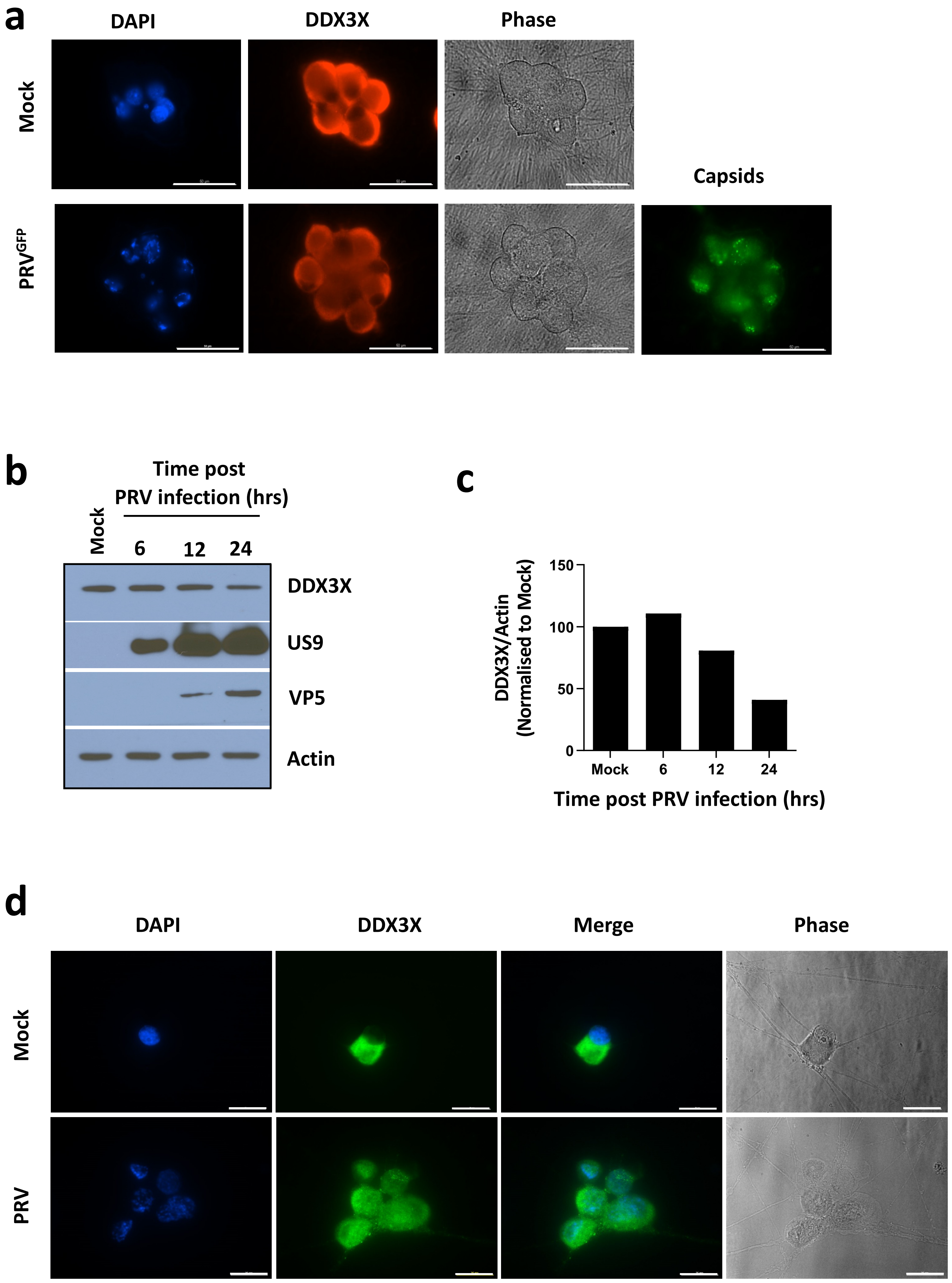

SUPPLEMENTAL FIGURE 1

**Supplemental Figure 1. PRV infects DRG neurons and reduces DDX3X levels.**

**a,** Representative immunofluorescence images of DDX3X levels in cultured SCG neurons mock infected and infected with GFP-labelled PRV for 12 hours. DAPI and phase contrast are also shown. Productive infection by PRV was confirmed by viral capsid staining. Scale bar, 50µm. **b,c,** Western blot time course of dorsal root ganglion (DRG) neurons infected with PRV and blotted for DDX3X and the PRV proteins US9 and VP5, assayed 6, 12 and 24 hours post infection. Actin is used as a loading control (**b**). Quantification of DDX3X levels normalized to mock treatment (no virus infection) (**c**). **d,** Representative immunofluorescence images of DDX3X levels in cultured DRG neurons mock infected and infected with PRV for 12 hours. DAPI and phase contrast is also shown. Scale bar, 25µm.

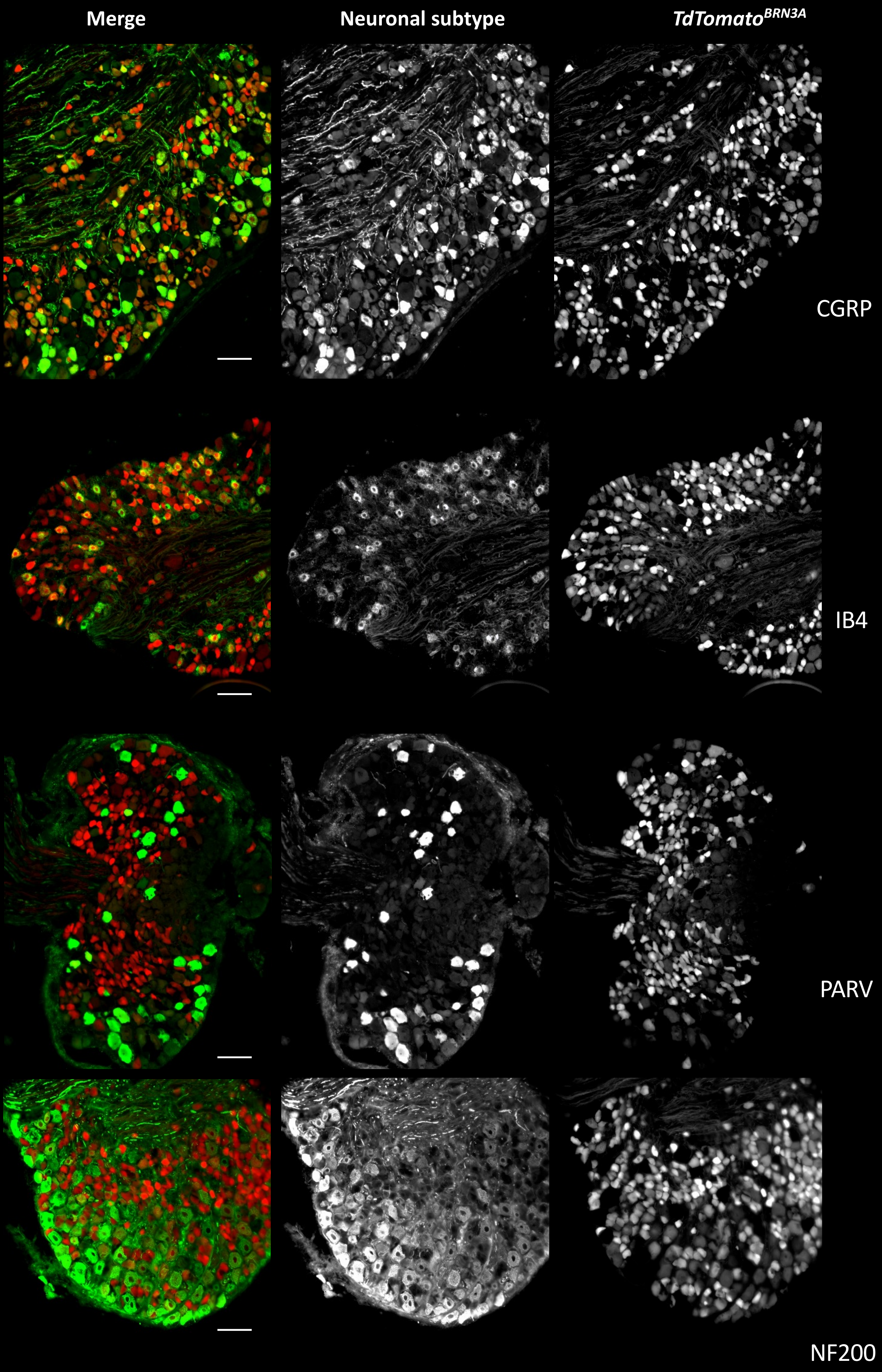

SUPPLEMENTAL FIGURE 2

**Supplemental Figure 2. Specific targeting of tamoxifen inducible *Brn3a-Cre* expression to small diameter sensory neurons.**

Representative images of DRG tissue from *TdTomato*$^{floxSTOPflox}$ reporter mice bred to the tamoxifen-inducible *Brn3A-Cre*$^{ERT}$ line to generate *TdTomato*$^{BRN3A}$ mice, analysed 2 weeks after tamoxifen treatment. Endogenous *TdTomato* marks the *Cre*-expressing neurons. Various neuronal subtypes were co-stained with CGRP (marks peptidergic, small-diameter sensory neurons), IB4-binding (marks non-peptidergic, small-diameter sensory neurons), Parv (marks large-diameter proprioceptors), and NF200 (marks large-diameter A-beta fibers) to check specificity of the *Brn3A-Cre* line. Merged images are also shown. Scale bar, 100µm.

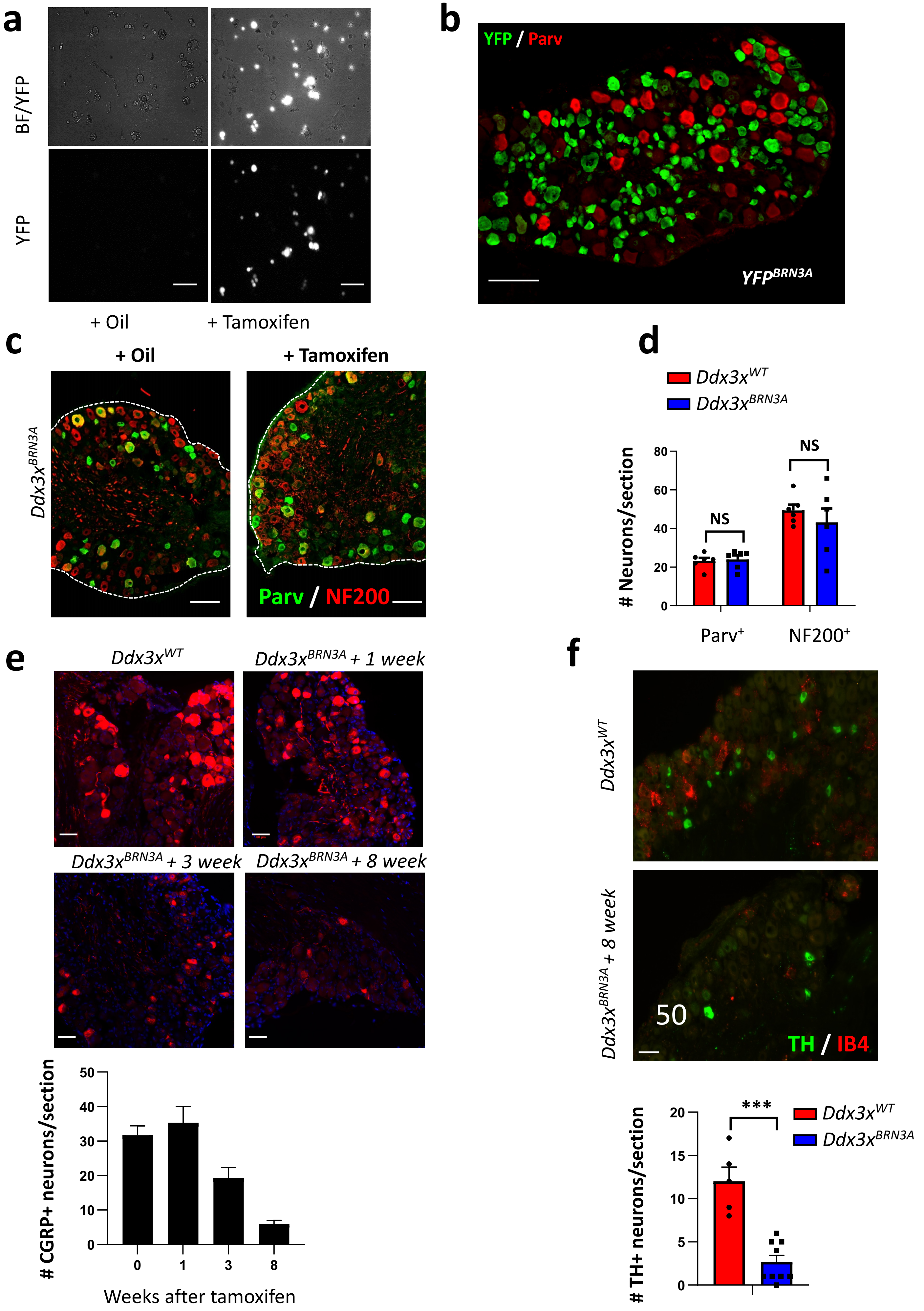

SUPPLEMENTAL FIGURE 3

**Supplemental Figure 3. Specific tamoxifen inducible *Brn3a-Cre* deletion in small diameter sensory neurons.**

**a,** Fluorescence and bright field (BF) images of cultured DRG neurons from $YFP^{BRN3A}$ mice 2 weeks after tamoxifen and vehicle (oil) treatment. Scale bar, 100µm. **b,** Representative immunofluorescence images of neurons in DRG tissue from $YFP^{BRN3A}$ mice stained with anti-PARV and labelled endogenous YFP. Scale bar, 100µm. **c,** Representative immunofluorescence images neurons in DRG tissue from $Ddx3x^{BRN3A}$ mice 6 weeks after treatment with tamoxifen or vehicle (oil) stained with anti-PARV and anti-NF200. **d**, Quantification of the number of each neuronal subtype as shown in (b) and (c). **e**, Representative immunofluorescence images, and quantification, of neurons in DRG tissue from $Ddx3x^{BRN3A}$ mice indicated weeks after treatment with tamoxifen stained with CGRP and DAPI. Scale bar, 50µm. **f,** Representative immunofluorescence images, and quantification, of tyrosine hydroxylase-positive ($TH^+$) neurons in DRG tissue from control and $Ddx3x^{BRN3A}$ mice ten weeks after treatment with tamoxifen stained with TH and IB4. Scale bar, 50µm.

Individual mice for each genotype are shown (d). Data are shown as means ± s.e.m. Multiple t-test (d); Two-tailed unpaired Student's t test (f). ***P < 0.001; NS, not significant.

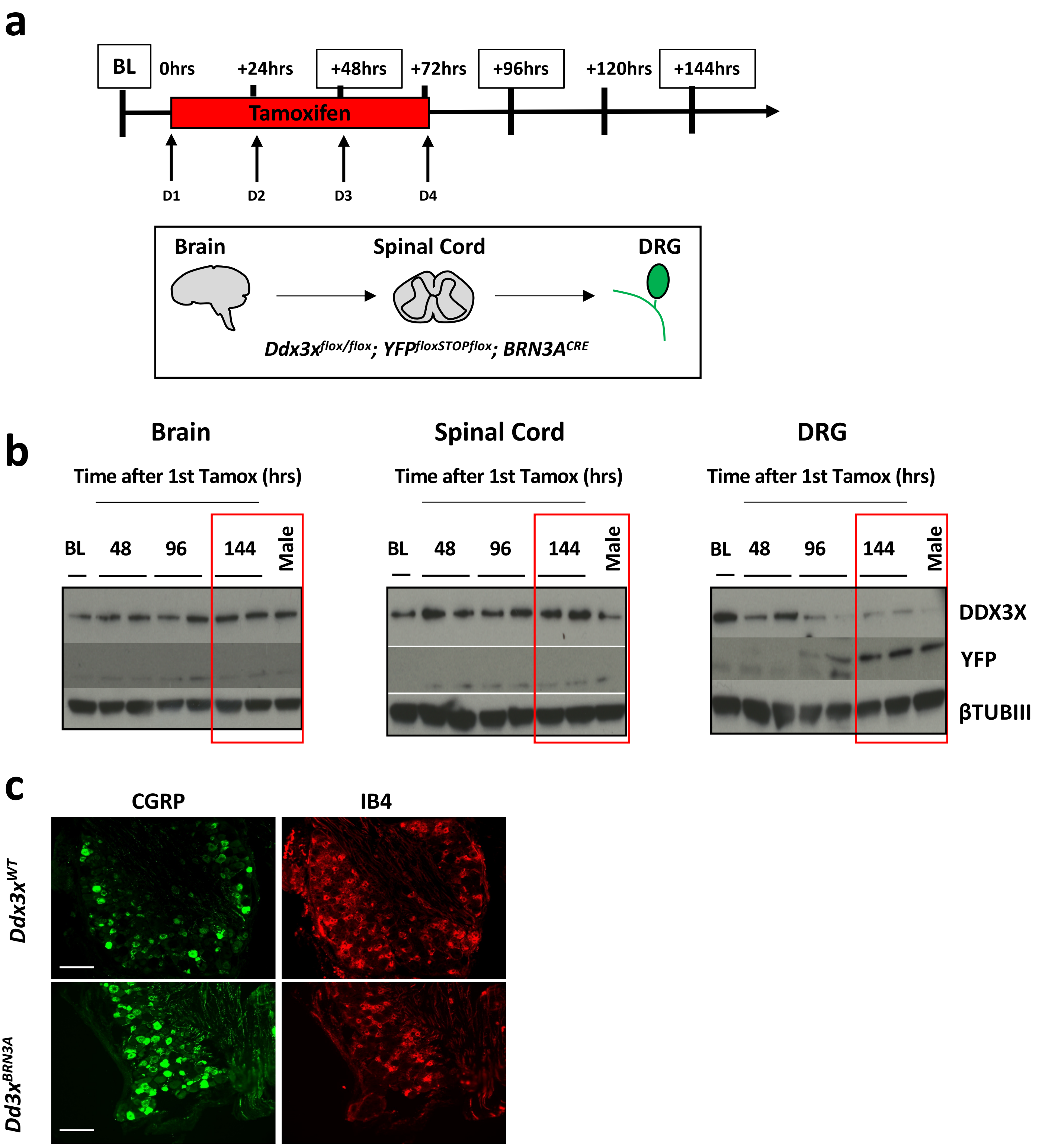

SUPPLEMENTAL FIGURE 4

**Supplemental Figure 4. Optimal time courses for RNA sequencing of DRG tissue of *Ddx3x*$^{WT}$ and *Ddx3x*$^{BRN3A}$ mice after tamoxifen treatment.**

**a,** Schematic to identify the optimal time point after tamoxifen treatment when *Brn3a-Cre* is expressed and DDX3X protein levels are diminished using the *Ddx3x*$^{flox/flox}$*; YFP*$^{floxSTOPflox}$*; BRN3A*$^{CRE}$ reporter line that only deletes in DRG neurons (green). **b**, Western blot analysis of YFP and DDX3X from the brain, spinal cord and DRG tissues to identify the optimal time point after tamoxifen treatment when DDX3X levels are diminished. Note the sensory neuronal specificity of the *Brn3A-Cre* line. **c**, Representative immunofluorescence images of neuronal subtypes in DRG tissue from *Ddx3x*$^{WT}$ and *Ddx3x*$^{BRN3A}$ mice 144 hours after initial tamoxifen treatment stained with anti-CGRP and IB4-lectin. Scale bar, 100µm.

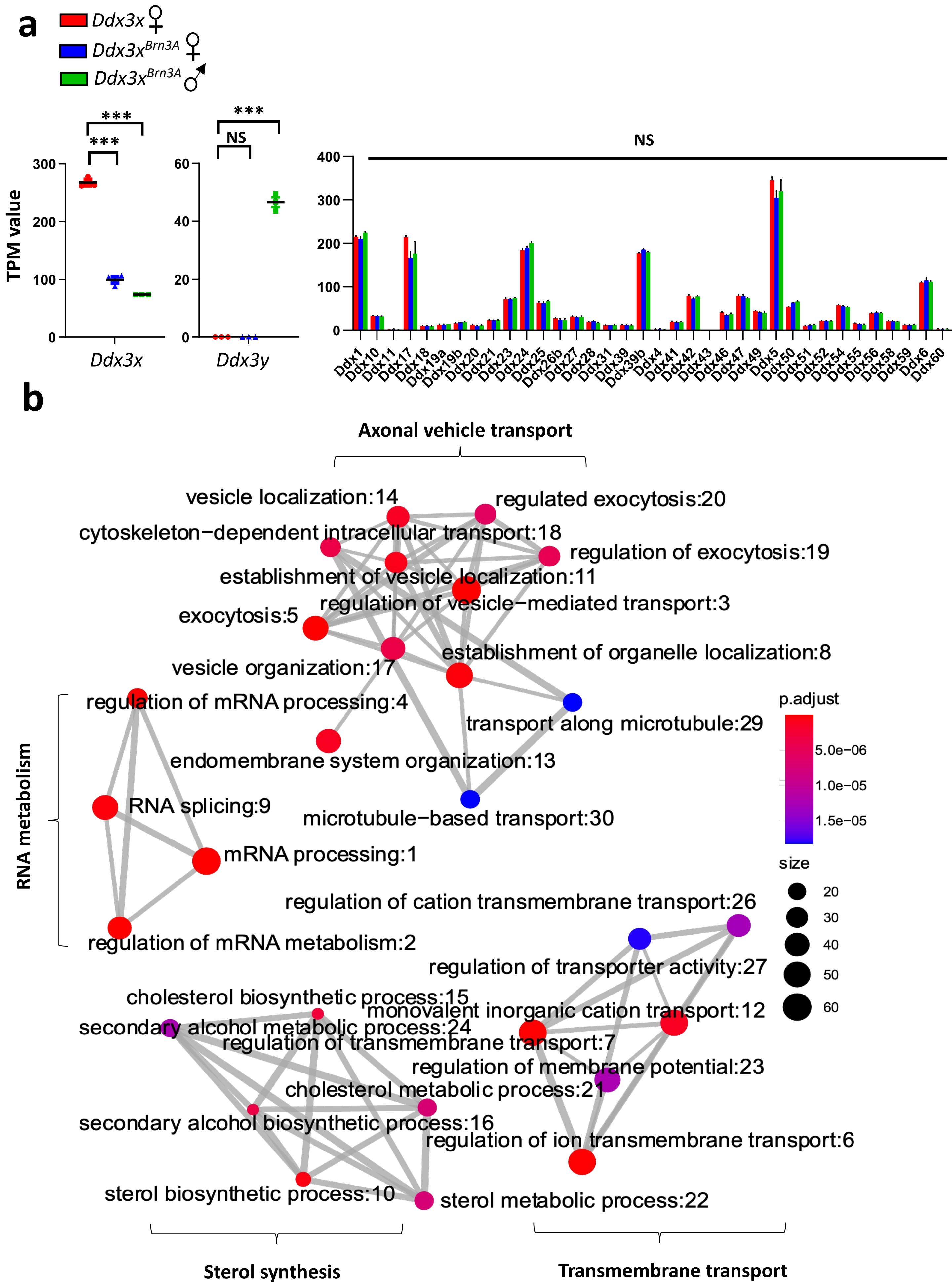

SUPPLEMENTAL FIGURE 5

**Supplemental Figure 5. Axonal and vesicle transport processes are de-regulated in response to sensory neuron-specific *Ddx3x* ablation.**

**a,** mRNA transcript levels of *Ddx3x* and *Ddx3y* (left panels) as well as all other *Ddx* family members from DRG tissue of female $Ddx3x^{WT}$ and $Ddx3x^{BRN3A}$ mice as well as male and $Ddx3x^{BRN3A}$ mice 144 hours after initial tamoxifen treatment. TPM, transcripts per million. **b,** Biological processes defined by gene ontology significantly downregulated upon *Ddx3x* deletion in sensory neurons from DRG tissue of female $Ddx3x^{WT}$ (N=3) and $Ddx3x^{BRN3A}$ (N=3) mice.

Individual mice for each genotype are shown (a). Scale bar, 100µm. Data are shown as means ± s.e.m. One-Way ANOVA with Dunnett's multiple comparison test (a). ***P < 0.001; NS, not significant.

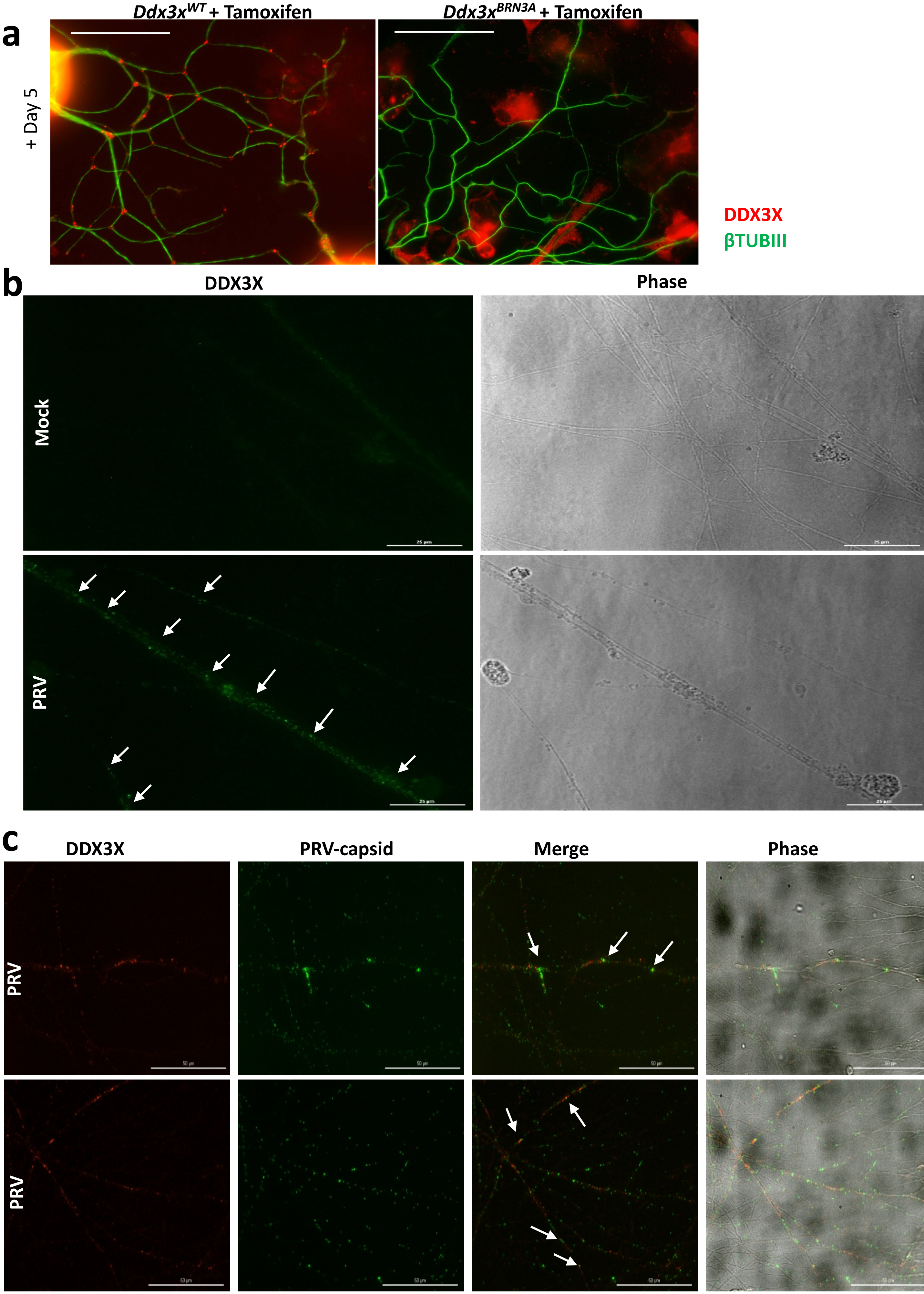

SUPPLEMENTAL FIGURE 6

**Supplemental Figure 6. PRV infection increases axonal DDX3X localization.**

**a**, Representative immunofluorescence images of DDX3X (red) and beta-Tubulin-III (green) expression in cultured DRG neurons from female *Ddx3x*$^{WT}$ and *Ddx3x*$^{BRN3A}$ mice 5 days after tamoxifen treatment. Note localization of DDX3X primarily at the branching points of the neurites of *Ddx3x*$^{WT}$ neurons. Scale bar, 50µm. **b,** Representative immunofluorescence images of cultured wild type DRG neurons 12 hours after infection with PRV, stained with anti-DDX3X (arrows). Phase contrast is also shown. Note increased DDX3X expression in PRV-infected neurites. Scale bar, 25µm. **c,** Representative immunofluorescence images of cultured wild type DRG neurons 12 hours after infection with GFP-capsid labelled PRV, stained with anti-DDX3X. Phase contrast is also shown. Arrows in the merged images indicate co-localization of DDX3X with PRV-capsid. Scale bar, 50µm.

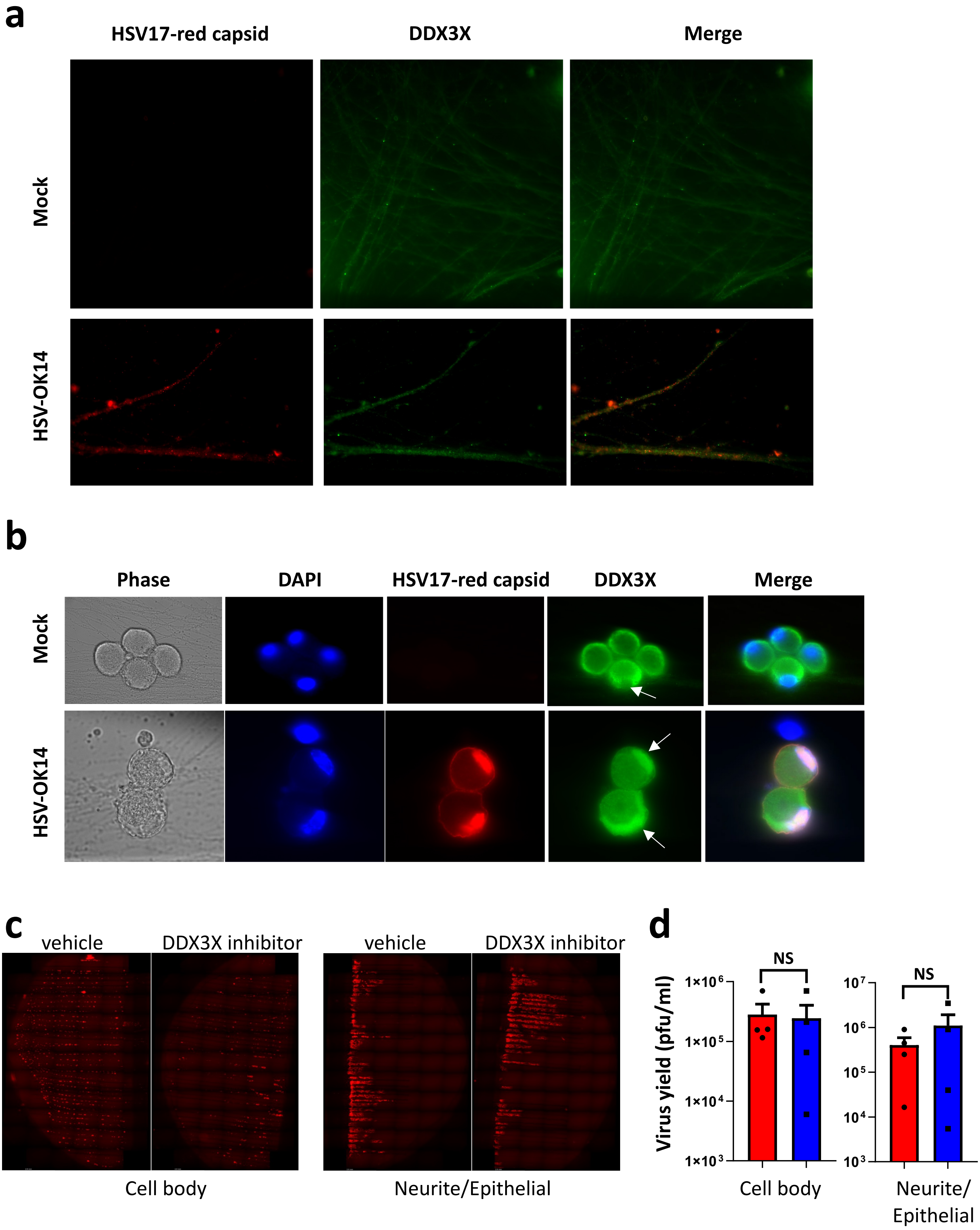

SUPPLEMENTAL FIGURE 7

**Supplemental Figure 7. DDX3X is not required for HSV-1 axonal transport.**

**a,** Representative immunofluorescence images of DDX3X in axons of cultured SCG neurons mock infected and infected with red capsid-labelled HSV-1 for 24 hours. **b,** Representative immunofluorescence images of DDX3X in cultured SCG neurons mock infected and infected with red capsid-labelled HSV-1 for 12 hours. DAPI and phase contrast are also shown. Productive infection by HSV-1 was confirmed by viral capsid staining. **c, d,** Representative immunofluorescence images of labelled-PRV in pig PK15 epithelial cells after cell body infection with HSV-1 in the tri-chamber system. Vehicle and DDX3X inhibitor were added one hour after PRV infection. Images were acquired at 24 hours post infection (**c**). Quantification of PRV yield from the cell body and neurite/epithelial compartments 24 hours post infection after vehicle and DDX3X inhibitor treatments (**d**).

Individual samples for each experiment are shown (d). Data are shown as means ± s.e.m. Two-tailed unpaired Student's t test (d). $^{**}P < 0.01$; NS, not significant.

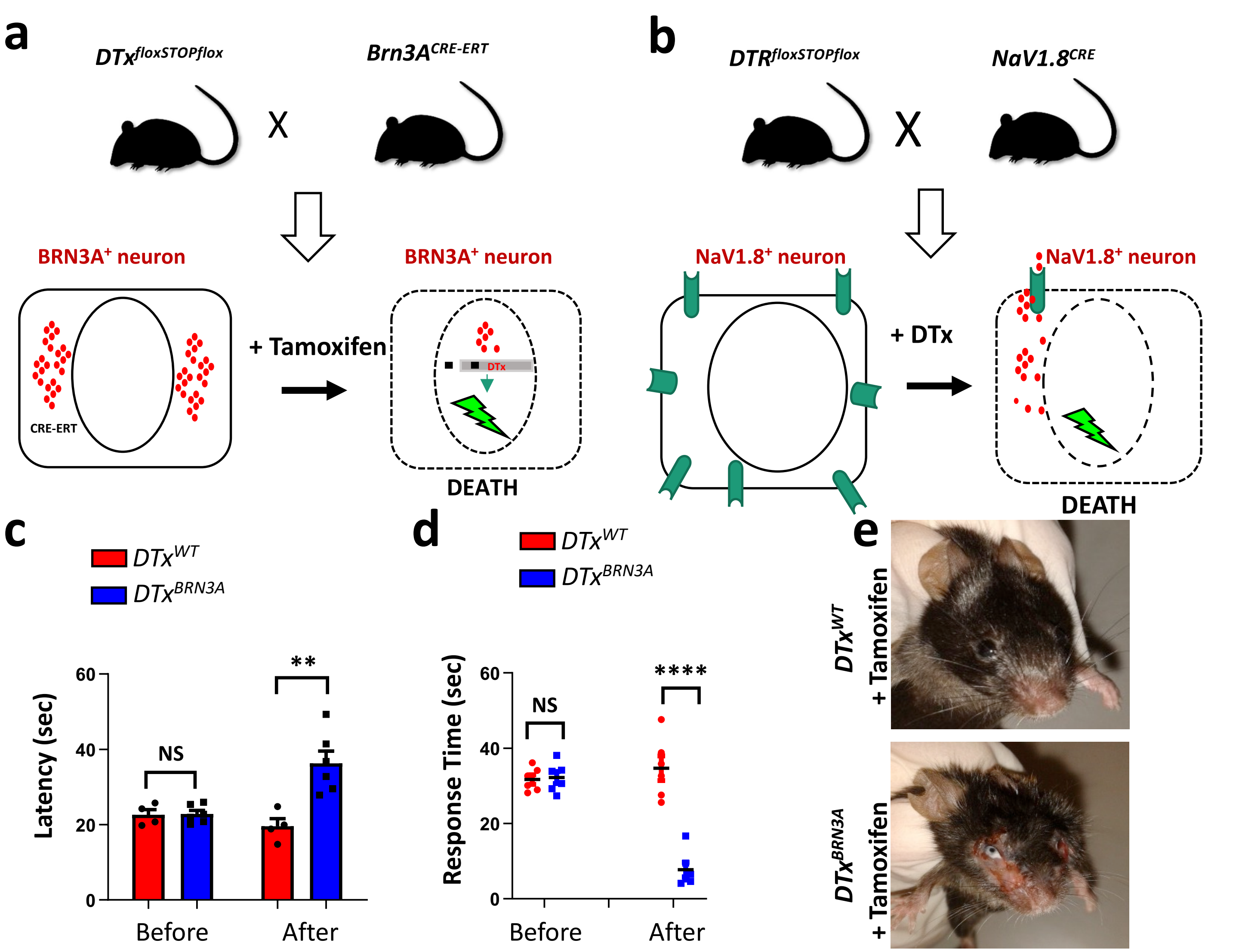

f

| | Lesion incidence |
|---|---|
| *Ddx3x^BRN3A* - Females | 8/8 |
| *Ddx3x^BRN3A* - Males | 0/4 |
| DTx^BRN3A - Females | 3/3 |
| DTx^BRN3A - Males | 3/3 |
| DTx^WT - Females | 0/4 |
| DTx^WT - Males | 0/2 |

g

| | | | Lesion incidence |
|---|---|---|---|
| >8 weeks | Male | [DTR^NaV1.8] + DTx | 7/7 |
| | Female | [DTR^NaV1.8] + DTx | 10/10 |
| 3 weeks | Male | [DTR^NaV1.8] + DTx | 0/3 |
| | | [DTR^WT] + DTx | 0/3 |
| | | [WT^NaV1.8] + DTx | 0/3 |
| | Female | [DTR^NaV1.8] + DTx | 0/3 |
| | | [DTR^WT] + DTx | 0/4 |
| | | [WT^NaV1.8] + DTx | 0/3 |

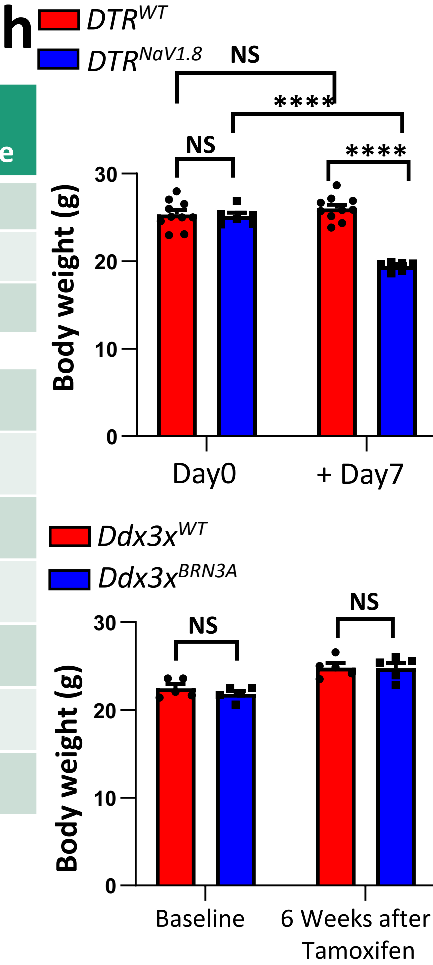

SUPPLEMENTAL FIGURE 8

**Supplemental Figure 8. Generation of genetic lines to abolish small-diameter sensory neurons.**

**a,** Schematic depicting the tamoxifen-inducible expression of diphtheria toxin (DTx) to cause death in *Brn3A+* sensory neurons. **b,** Schematics depicting the inducible ablation of *NaV1.8+* sensory neurons by the administration of DTx to $DTR^{NaV1.8}$ mice in which the diphtheria toxin receptor (DTR) is selectively expressed by *NaV1.8+* sensory neurons. **c**, Contact heat withdrawal latencies using the hotplate assay at 52°C for control $DTx^{WT}$ (N=4) and littermate $DTx^{BRN3A}$ (N=6) mice before and two weeks after tamoxifen treatment. **d,** Nocifensive responses in a 3-minute time frame after intraplantar injection of 2 µg of capsaicin into the left hind paw of control $DTx^{WT}$ (N=6) and littermate $DTx^{BRN3A}$ (N=6) mice before and two weeks after tamoxifen treatment. **e,** Representative images of skin lesions on the face of $DTx^{BRN3A}$ mice housed separately 3 weeks after tamoxifen treatment. A littermate control $DTx^{WT}$ mouse is shown as control. **f,g,** Prevalence of skin lesions in various mouse lines and genders treated with tamoxifen (**f**) and in various DTR lines treated with DTx (**g**). **h,** Body weight measurements in $DTR^{NaV1.8}$ mice treated with DTx (upper panel) and $Ddx3x^{BRN3A}$ female mice treated with tamoxifen (lower panel).

Individual mice for each genotype are shown (c,d,h). Data are shown as means ± s.e.m. Multiple t-test (c,d); Two-way ANOVA with Tukey's multiple comparison test (h). $^{**}P < 0.01$; $^{****}P < 0.0001$; NS, not significant.

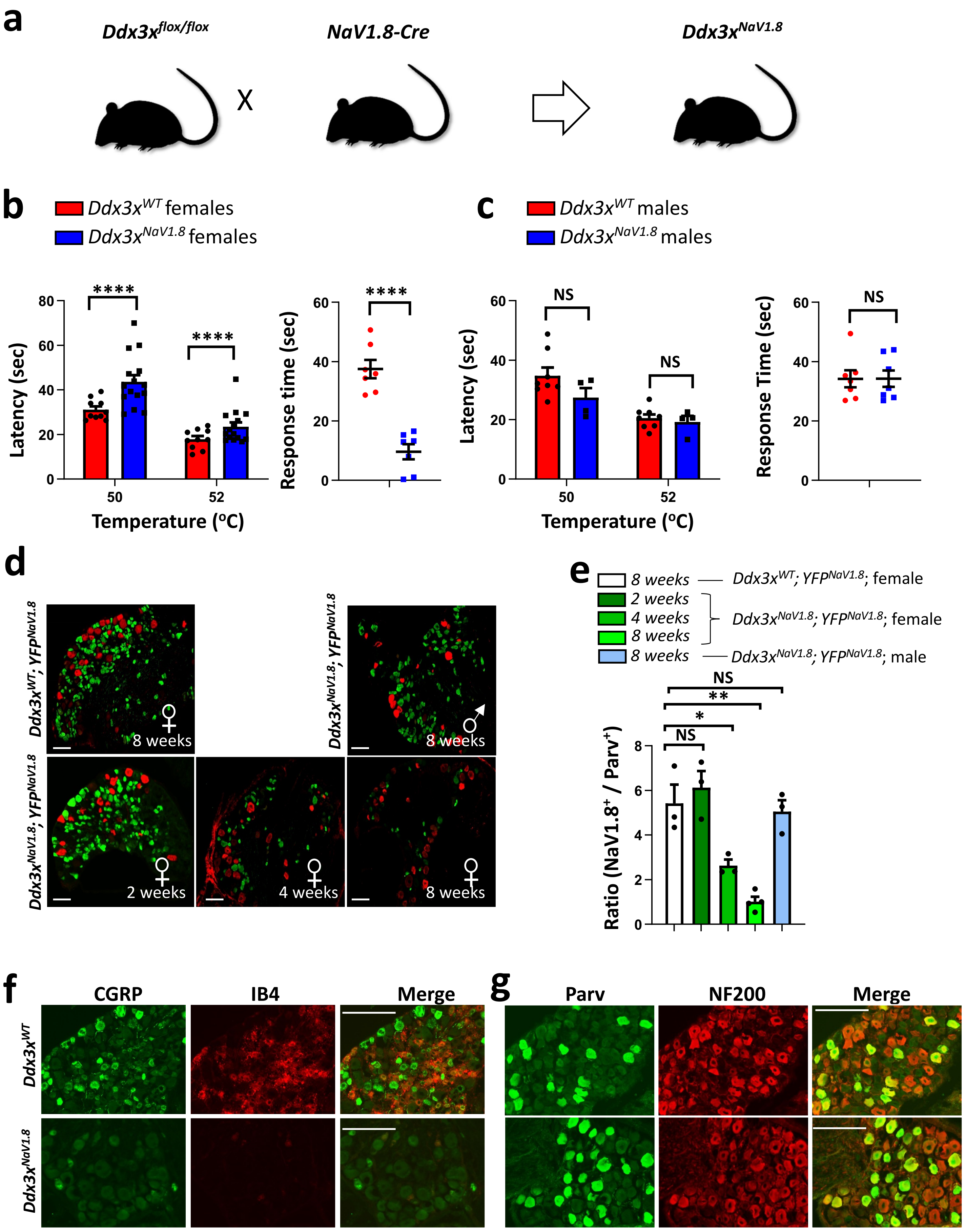

SUPPLEMENTAL FIGURE 9

**Supplemental Figure 9. Developmental loss of *Ddx3x* does not lead to "mad itch".**

**a,** Schematic breeding to developmentally ablate Ddx3x in small diameter peripheral sensory neurons using the *NaV1.8-Cre* line. **b**, Contact heat withdrawal latencies using the hotplate assay at the indicated temperatures in 12-week old control $Ddx3x^{WT}$ (N=10) and littermate $Ddx3x^{NaV1.8}$ (N=15) female mice (left panel). Nocifensive responses in a 3-minute time frame after intraplantar injection of 2 µg of capsaicin into the left hind paw of control $Ddx3x^{WT}$ (N=7) and littermate $Ddx3x^{NaV1.8}$ (N=7) female mice (right panel). Experiment was repeated an additional time with similar results. **c,** Contact heat withdrawal latencies using the hotplate assay at the indicated temperatures for 12-week old control $Ddx3x^{WT}$ (N=8) and littermate $Ddx3x^{NaV1.8}$ (N=4) male mice (left panel). Nocifensive responses in a 3-minute time frame after intraplantar injection of 2 µg of capsaicin into the left hind paw of control $Ddx3x^{WT}$ (N=7) and littermate $Ddx3x^{NaV1.8}$ (N=7) male mice (right panel). Experiment was repeated an additional time with similar results. **d,** Representative immunofluorescence images of neurons in DRG tissue from control $Ddx3x^{WT}$; $YFP^{NaV1.8}$ and $Ddx3x^{NaV1.8}$; $YFP^{NaV1.8}$ mice at the indicated ages and sex stained with anti-PARV (red) and endogenous YFP (green). Scale bar, 100µm. **e,** Quantification of neuronal subtypes from (d). **f,g,** Representative immunofluorescence images of neurons in DRG tissue from 12-week old control $Ddx3x^{WT}$ and $Ddx3x^{NaV1.8}$ female mice stained with the indicated markers to delineate neuronal subtypes. Scale bar, 100µm.

Individual mice for each genotype are shown (b,c,e). Data are shown as means ± s.e.m. Multiple t-test (b,c left panels); Two-tailed unpaired Student's t test (b,c right panels); One-way ANOVA with Dunnett's multiple comparison test (e). *P < 0.05; **P < 0.01; ****P < 0.0001; NS, not significant.

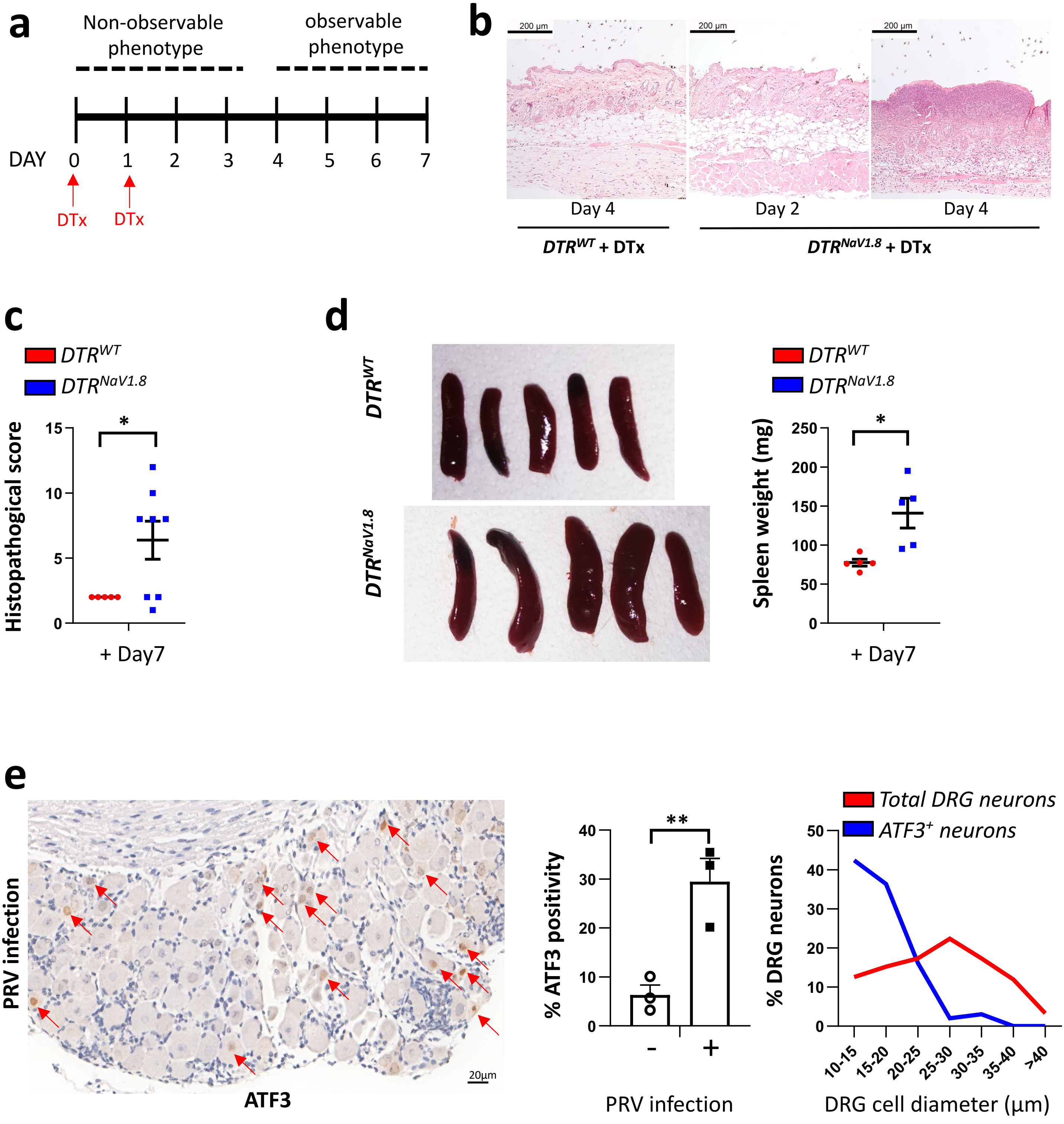

SUPPLEMENTAL FIGURE 10

**Supplemental Figure 10. DTx-treated *DTR^{NaV1.8}* mice develop inflammatory skin lesions.**

**a,** Schematic showing the time course of skin lesions in *DTR^{NaV1.8}* mice treated twice with DTx (indicated with red arrows). **b,** Representative H&E histology sections of nape skin from 8-week-old *DTR^{WT}* and littermate *DTR^{NaV1.8}* mice at the indicated days after DTx treatment. **c**, Histopathological scoring (see methods) of nape skin sections of 8-week-old *DTR^{WT}* and littermate *DTR^{NaV1.8}* mice 7 days after DTx treatment. **d,** Left: macroscopic images of spleens from 8-week-old *DTR^{WT}* and *DTR^{NaV1.8}* mice 7 days after DTx treatment. Right: quantification of spleen weights. **e**, Representative histological image of ATF-3 staining (left panel) and quantification (middle panel) and neuron-diameter frequency of ATF-$3^+$ neurons (right panel) of ATF-3-positive DRG neurons from N=3 wild type 7 hours post infection with PRV.

Individual mice for each genotype are shown (c,d). Data are shown as means ± s.e.m. Two-tailed unpaired Student's t test (c,d). *P < 0.0; NS, not significant.

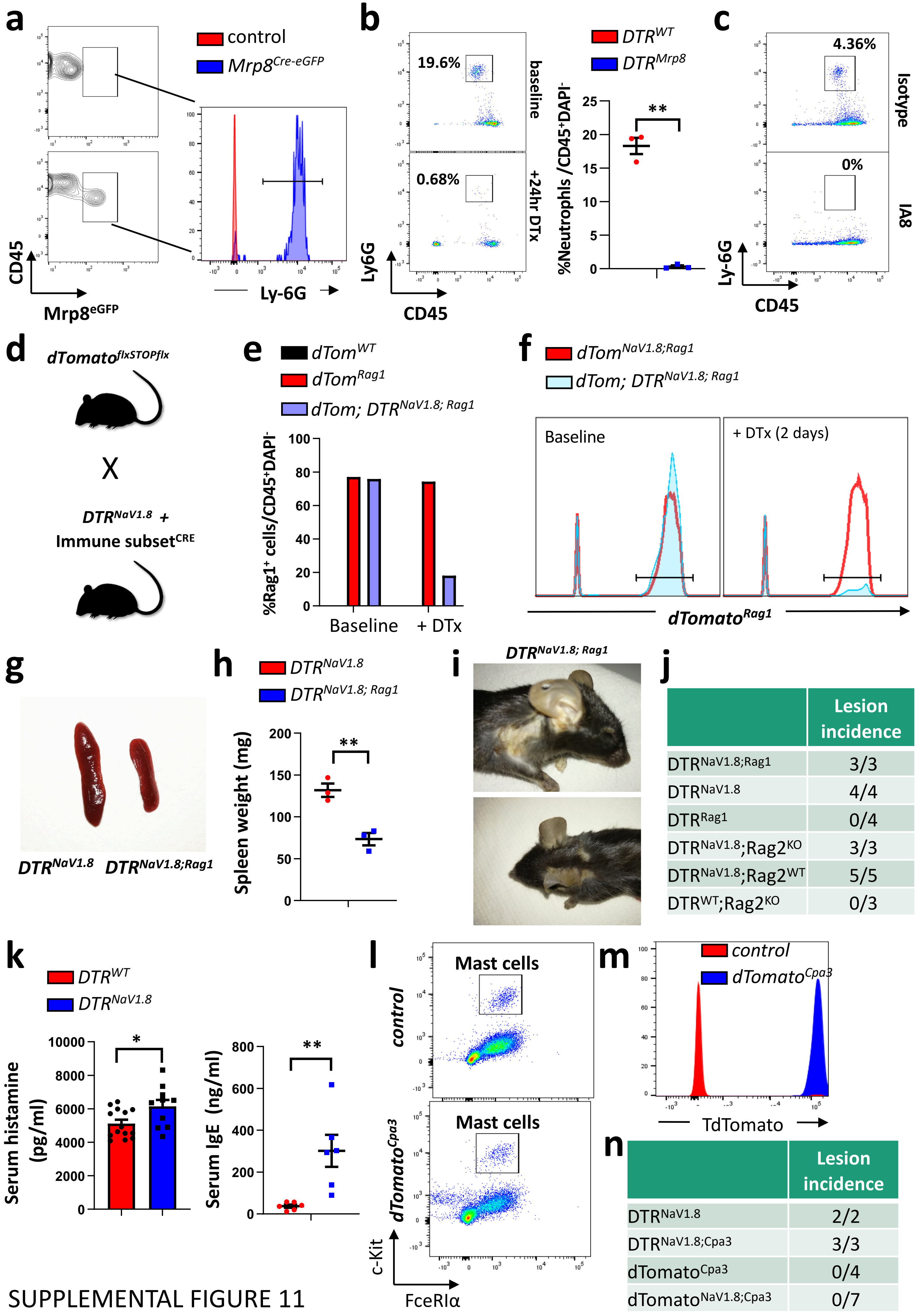

j

| | Lesion incidence |
|---|---|
| DTR$^{NaV1.8;Rag1}$ | 3/3 |
| DTR$^{NaV1.8}$ | 4/4 |
| DTR$^{Rag1}$ | 0/4 |
| DTR$^{NaV1.8}$;Rag2$^{KO}$ | 3/3 |
| DTR$^{NaV1.8}$;Rag2$^{WT}$ | 5/5 |
| DTR$^{WT}$;Rag2$^{KO}$ | 0/3 |

n

| | Lesion incidence |
|---|---|
| DTR$^{NaV1.8}$ | 2/2 |
| DTR$^{NaV1.8;Cpa3}$ | 3/3 |
| dTomato$^{Cpa3}$ | 0/4 |
| dTomato$^{NaV1.8;Cpa3}$ | 0/7 |

SUPPLEMENTAL FIGURE 11

**Supplemental Figure 11. No obvious involvement of prominent immune cells in the prevalence of "mad itch".**

**a,** Use of the *Mrp8*$^{Cre\text{-}eGFP}$ mice to specifically delete neutrophils (stained with anti-Ly6G) in our "mad itch" model. **b**, Left: Percentages of Ly6G$^{+}$ blood neutrophils among total CD45$^{+}$ immune cells in *DTR*$^{Mrp8}$ mice at baseline and 2 days after DTx treatment. Right: quantification of the percentages of neutrophils in total CD45$^{+}$ blood cells in 8-week-old *DTR*$^{WT}$ (N=3) and *DTR*$^{Mrp8}$ (N=3) mice, 7 days after DTx treatment. **c**, Efficiency of splenic LyG6+ neutrophil depletion after two days of saline treatment or 1A8 (anti-Ly-6G) depleting antibody (20 µg, i.p. administration per mouse) treatment in wild type animals. **d,** Breeding scheme to deplete immune cell types with specific *Cre*-expressing transgenic lines in addition to labelling the immune cell type of interest (*dTomato*$^{floxSTOPflox}$) and ablating sensory neurons (*DTR*$^{NaV1.8}$) upon DTx treatment. **e,f**, Depletion efficiency of T and B cells using the *Rag1*-expressing *Cre* line in conjunction with *dTomato*$^{NaV1.8;\ Rag1}$ mice after 2 days of DTx treatment. **g,h,** Representative images (**g**) and weights (**h**) of spleens from *DTR*$^{NaV1.8}$ (N=3) and *DTR*$^{NaV1.8;Rag1}$ (N=3) mice after 7 days of DTx treatment. **i,j,** Representative images of skin lesions on the face, flank and nape of *DTR*$^{NaV1.8;Rag1}$ mice housed separately 7 days after DTx treatment (**i**) and the prevalence of skin lesions in various mouse lines with T and B lymphocyte ablations (**j**). **k**, Serum histamine and IgE levels from 8-week-old *DTR*$^{WT}$ (N=14 and 6, respectively) and *DTR*$^{NaV1.8}$ (N=10 and 6, respectively) mice 7 days after DTx treatment. **l,m**, Use of the *dTomato* reporter line crossed to the mast cell and basophil specific *Cpa3-Cre* line. **n**, Prevalence of self-inflicted skin lesions in the sensory neuron- and mast cell/basophil cell-ablated *DTR*$^{NaV1.8;Cpa3}$ line, assessed 7 days after DTx treatment.

Individual mice for each genotype are shown (b,h,k). Data are shown as means ± s.e.m. Two-tailed unpaired Student's t test (b,h,k). *P < 0.05; **P < 0.01.

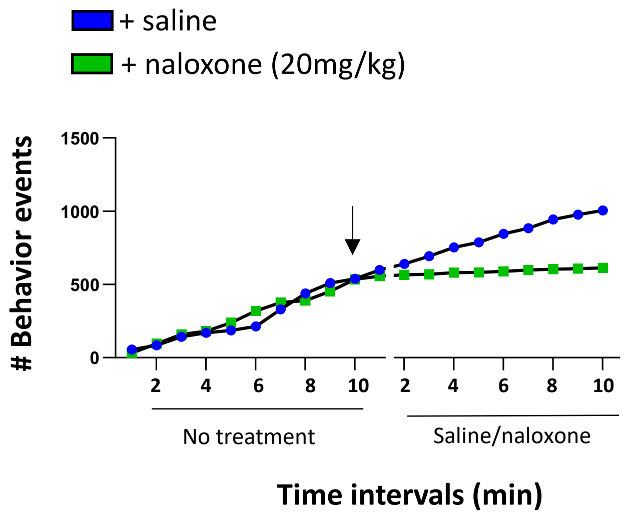

SUPPLEMENTAL FIGURE 12

**Supplemental Figure 12. Naloxone treatment halts the "mad itch".**

Effect of saline and naloxone treatment (20mg/kg, i.p. administration) on "mad itch" behavioral events in a representative *Ddx3x*$^{BRN3A}$ mouse 6 weeks after tamoxifen treatment.

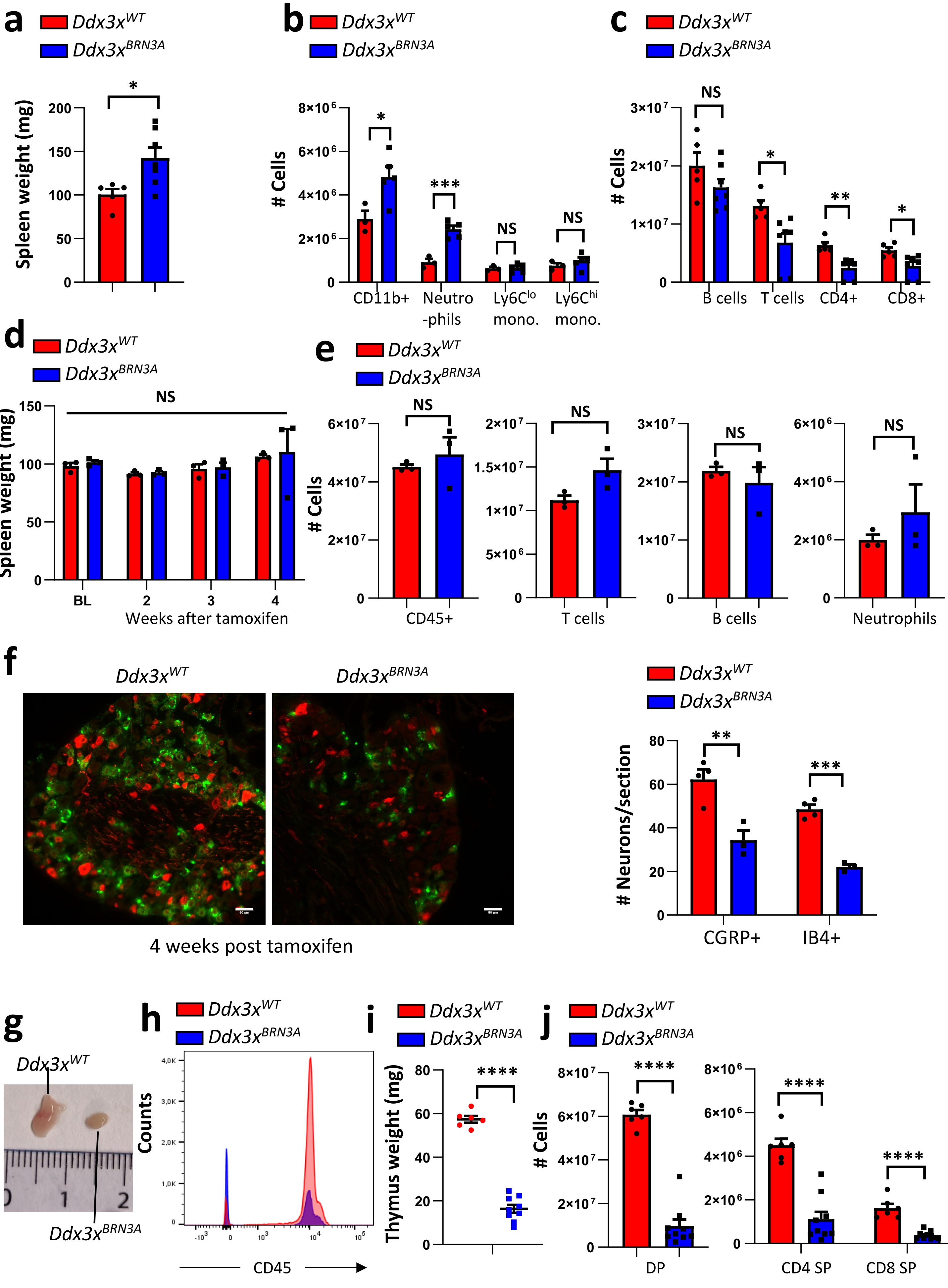

SUPPLEMENTAL FIGURE 13

**Supplemental Figure 13. Progressive immune deviations upon *Ddx3x* deletion in sensory neurons.**

**a**, Spleen weights from 14-week-old *Ddx3x*$^{WT}$ (N=5) and littermate *Ddx3x*$^{BRN3A}$ (N=7) mice 6 weeks after tamoxifen treatment. **b,c,** Number of the various indicated cell types in the spleen of 14-week-old *Ddx3x*$^{WT}$ (N=5) and littermate *Ddx3x*$^{BRN3A}$ (N=7) mice 6 weeks after tamoxifen treatment. **d,** Spleen weights of control *Ddx3x*$^{WT}$ (N=3) and littermate *Ddx3x*$^{BRN3A}$ (N=3) female mice at the indicated time points after tamoxifen treatment. e**,** Cell numbers of all haematopietic cells (CD45$^{+}$) and the indicated splenic cellular subpopulations of control *Ddx3x*$^{WT}$ (N=3) and littermate *Ddx3x*$^{BRN3A}$ (N=3) female mice at 4 weeks after tamoxifen treatment. **f,** Representative immunofluorescence images of CGRP$^{+}$ and IB4-binding neurons within DRG tissue and quantification of CGRP$^{+}$ and IB4$^{+}$ neuronal numbers/section from *Ddx3x*$^{WT}$ and *Ddx3x*$^{BRN3A}$ mice 4 weeks after tamoxifen treatment. Scale bar, 50µm. **g,** Representative image of thymi from 14-week-old *Ddx3x*$^{WT}$ and littermate *Ddx3x*$^{BRN3A}$ mice, assayed 6 weeks after tamoxifen treatment. **h,i,** Representative FACS histograms of CD45$^{+}$ thymocytes gated on viable cells (**h**) and thymus weights (**i**) of 14-week-old *Ddx3x*$^{WT}$ (N=6) and littermate *Ddx3x*$^{BRN3A}$ (N=9) mice 6 weeks after tamoxifen treatment. **j**, Number of double positive CD4$^{+}$CD8$^{+}$ (DP) and CD4$^{+}$ or CD8$^{+}$ single positive (SP) cells in the thymus of 14-week-old *Ddx3x*$^{WT}$ (N=6) and littermate *Ddx3x*$^{BRN3A}$ (N=9) mice, analyzed 6 weeks after tamoxifen treatment.

Individual mice for each genotype are shown (a-f, i,j). Data are shown as means ± s.e.m. Two-tailed unpaired Student's t test (a,e,I,j); Multiple t-test (b,c,d,j);. *P < 0.05; **P < 0.01; ***P < 0.001; ****P < 0.0001; NS, not significant.

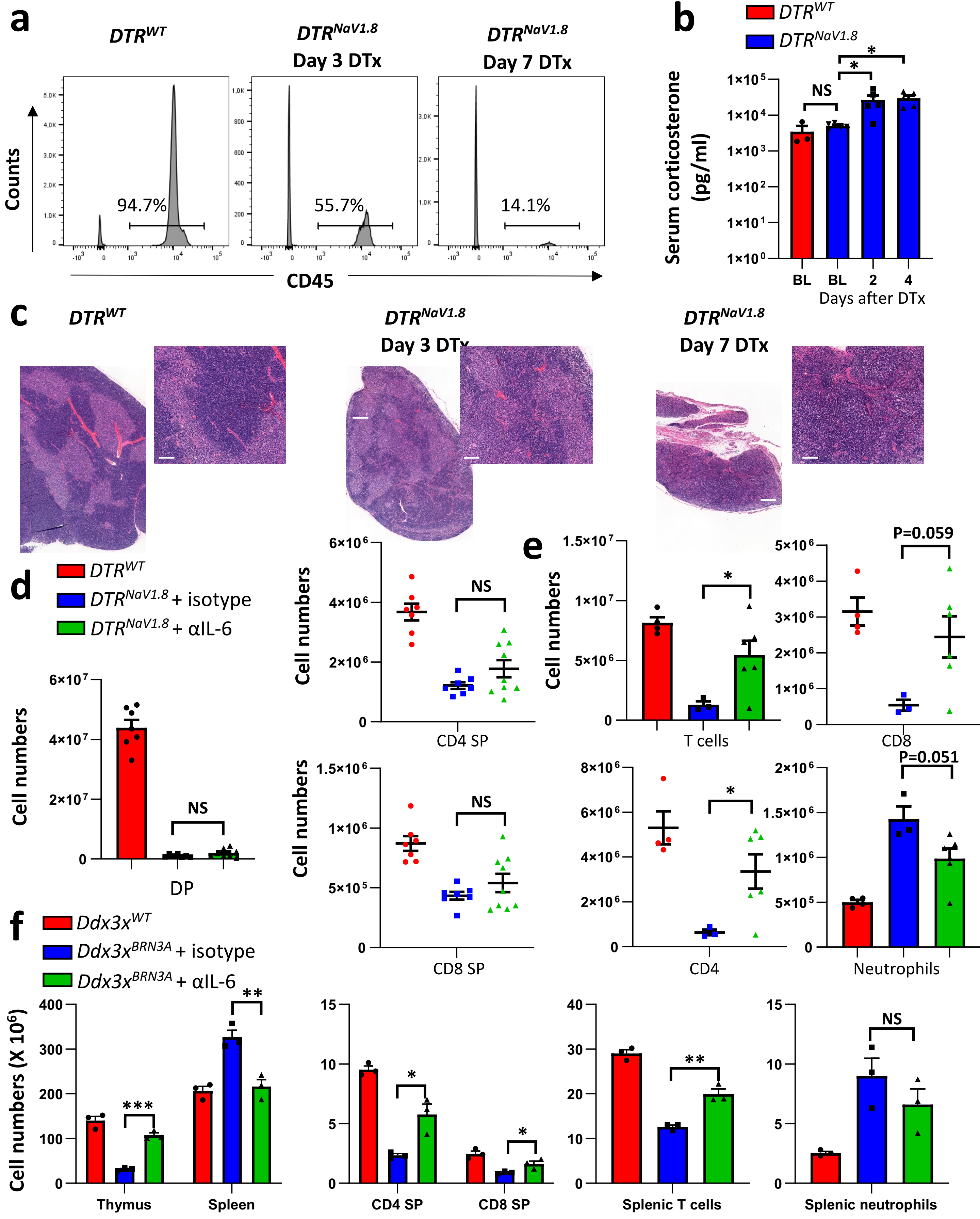

**g**

| | Lesion incidence | | Lesion incidence |
|---|---|---|---|
| DTR$^{NaV1.8}$ + isotype | 6/6 | DTR$^{NaV1.8}$ + αIL-6 | 6/6 |
| *Ddx3x$^{BRN3A}$* + isotype | 3/3 | *Ddx3x$^{BRN3A}$* + αIL-6 | 3/3 |

SUPPLEMENTAL FIGURE 14

**Supplemental Figure 14. Sensory neuronal loss induces thymic atrophy.**

**a**, Representative FACS histograms of $CD45^+$ thymocytes from *$DTR^{WT}$* and littermate *$DTR^{NaV1.8}$* mice 3 and 7 days after DTx treatment. **b**, Serum corticosterone levels in control *$Ddx3x^{WT}$* (N=3) mice at baseline and in littermate *$DTR^{NaV1.8}$* (N=5) mice at baseline and days indicated after DTx administration. **c**, Representative H&E stainings of the thymi from *$DTR^{WT}$* and littermate *$DTR^{NaV1.8}$* mice 3 and 7 days after DTx treatment. **d,** Numbers of CD4+CD8+ double positive (DP), CD4+ single positive (SP) and CD8+ SP thymocytes control *$DTR^{WT}$* (N=7), *$DTR^{NaV1.8}$* treated with isotype control antibodies (N=7) and *$DTR^{NaV1.8}$* treated with 100µg (every day starting one day after first DTx administration) anti-IL-6 antibodies (N=9), 7 days after DTx treatment. Individual mice for each genotype are shown. **e**, Numbers of total $TCR\alpha\beta^+$ T cells, $CD4^+$ T cells, $CD8^+$ T cells, and $Ly6G^+$ neutrophils in the spleens of control *$DTR^{WT}$* mice (N=4), *$DTR^{NaV1.8}$* mice treated with isotype control antibody (N=3) and *$DTR^{NaV1.8}$* mice treated with 100µg (every day starting one day after first DTx administration) anti-IL-6 antibody (N=6) 7 days after DTx treatment. **f,** Numbers of total $TCR\alpha\beta^+$ T cells, $CD4^+$ T cells, $CD8^+$ T cells, and $Ly6G^+$ neutrophils in the spleens of control *$Ddx3x^{WT}$* (N=3), *$Ddx3x^{BRN3A}$* treated with isotype control antibody (N=3) and *$Ddx3x^{BRN3A}$* treated with 100µg (for 5 consecutive days one week post tamoxifen) anti-IL-6 antibody (N=3) 7 weeks after tamoxifen treatment. **g**, Table depicting the incidence of skin lesions in DTx-treated *$DTR^{NaV1.8}$* and tamoxifen-treated *$Ddx3x^{BRN3A}$* mice administered with isotype control of anti-IL-6 antibody as described above.

Individual mice for each genotype are shown (b,d-f). Data are shown as means ± s.e.m. One-way ANOVA with Dunnett's multiple comparison test (b). Two-tailed unpaired Student's t test (d-f). $^*P < 0.05$; $^{**}P < 0.01$; NS, not significant.

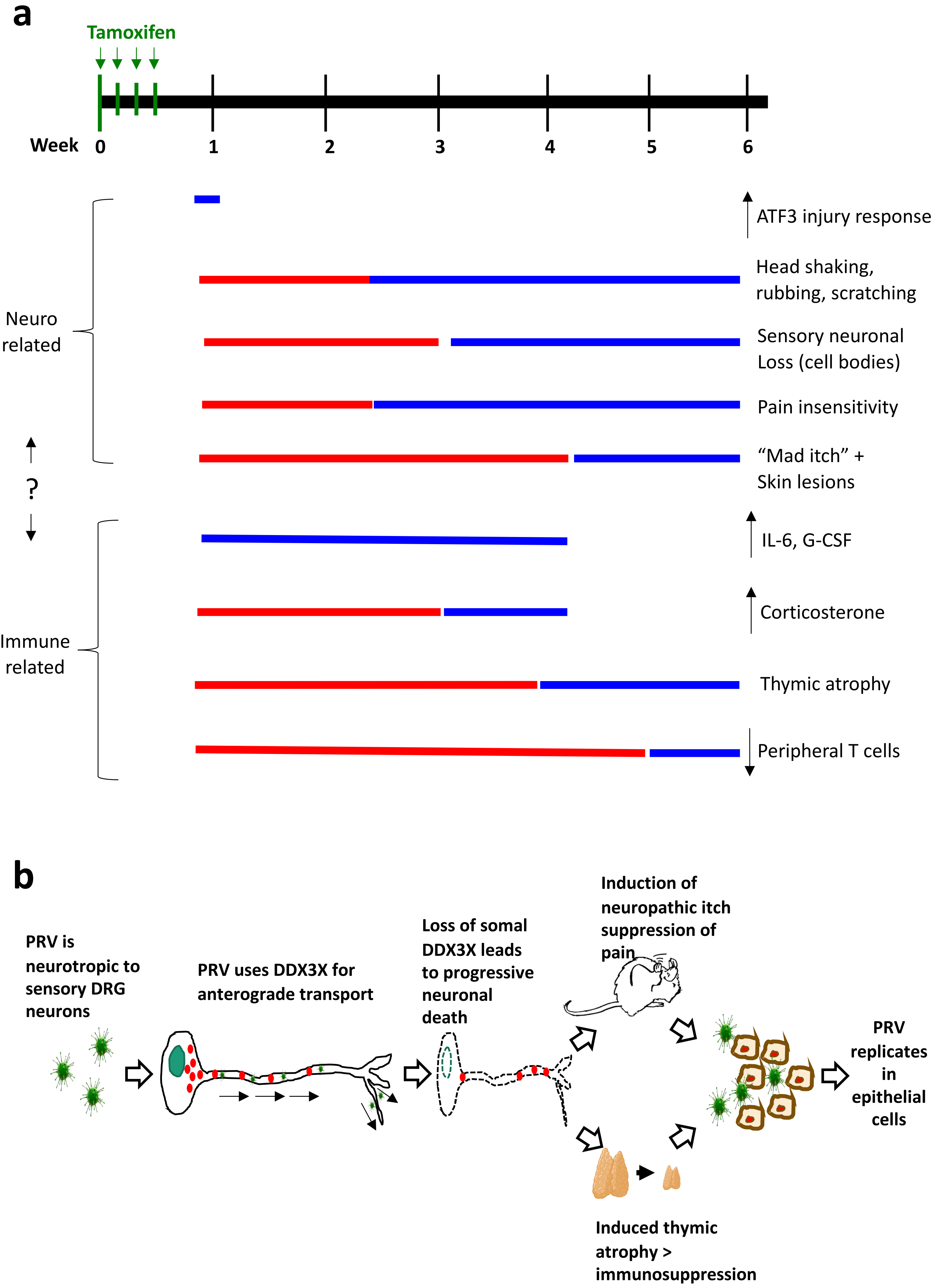

SUPPLEMENTAL FIGURE 15

**Supplemental Figure 15. Host-viral interactions involved in PRV spread and infectivity.**

**a,** Graphic depicting the time frames of the various neuro-immune pathophysiological effects caused by *Ddx3x* deletion in sensory neurons after tamoxifen treatment. **b,** Schematic model how PRV, by hijacking DDX3X, induces a series of pathophysiological mechanisms to aid infectivity and spread.

**Supplemental Table S1.**

| | Genetic specific *Ddx3x* deletion | PRV infection | References (PMID) | Animals |
|---|---|---|---|---|
| Reduced somal DDX3X levels | Y | Y | | mouse |
| Increase of serum IL-6 and G-CSF | Y | Y | 30258005 | mouse |
| "Mad itch" | Y | Y | [illegible]2759704, 12686430, 1314296[illegible] | cattle, dog, wolf, goat, sheep, fox, mink, raccoon |
| Sensory neuronal loss | Y | Y | [illegible]6326038; 11119615; 1732534[illegible] | pig, mouse |
| Reduced pain responses | Y | ND | | |
| Corticosterone increase | Y | ND | | |
| Injured DRG signature - increased ATF3 | Y | Y | | |
| Thymic atrophy | Y | Y | 16423576 | pig |
| Effect on peripheral immunity | Y | ND | | |
| Self-inflicted deep tissue lesions | Y | Y | [illegible]2759704, 12686430, 1314296[illegible] | cattle, dog, wolf, goat, sheep, fox, mink, raccoon |

### Supplemental Table

**Supplemental Table S1. Common characteristics shared by PRV infection in animals and sensory neuronal ablation of *Ddx3x*.**

Common pathological effects between PRV infection and sensory neuron specific temporal loss of *Ddx3x*. Y, yes (in common); ND, not determined. PMID, Pubmed ID.

### Supplemental Movies

**Supplemental Movie 1. “Mad itch”-like behaviors of a *Ddx3x$^{BRN3A}$* female mouse**.

Recording of a *Ddx3x$^{BRN3A}$* female mouse 6 weeks after tamoxifen administration. Note the self-inflicted lesions on the face and flanks/legs.

**Supplemental Movie 2. Typical “mad itch”-like behaviors of *Ddx3x$^{BRN3A}$* female mice**.

Recording of four *Ddx3x$^{BRN3A}$* female mice 6 weeks after tamoxifen administration.

**Supplemental Movies 3-6. The transport of PRV virions is diminished by inhibition of DDX3X.**

Representative time lapse microscopy of red labelled-PRV (PRV180-Red) infection of primary neurons in the tri-chamber system imaged in the cell body compartments (SV 3 and 4) and in the neurite/epithelial compartments (SV 5 and 6) images from 8 hours post infection each hour until 22 hours post infection. Samples were treated with vehicle (SV3 and 5) or DDX3X inhibitor (SV 4 and 6) into the cell body compartment at 1 hour post infection.
